# Vitamin C-Induced Photo-Redox Threshold Enables High-Fidelity Volumetric Printing of Pristine Collagen

**DOI:** 10.64898/2026.04.13.717972

**Authors:** Bin Wang, Amelia Hasenauer, Kristian Ivkovic, Ann-Sophie Frind, David Fercher, Marcy Zenobi-Wong

**Author notes:** B. Wang and A. Hasenauer contributed equally to this work.

## Abstract

Tomographic volumetric printing (TVP) enables rapid fabrication of complex, centimeter-scale 3D architectures. TVP of pristine proteins like collagen is attractive because it better preserves native bioactive motifs that regulate cell–matrix signaling. However, direct TVP of collagen remains challenging because dityrosine crosslinking, driven by visible-light-activated Ru(II)bpy_3_^2+^/sodium persulfate (SPS), lacks an effective inhibitory mechanism. This results in near-immediate crosslinking upon exposure to light, which leads to an insufficient nonlinear threshold response that fails to suppress background curing. Here, we introduce vitamin C (L-ascorbic acid) as a biocompatible redox regulator to overcome this limitation. UV–Vis kinetics demonstrate that vitamin C suppresses Ru(III) accumulation and scavenges persulfate radicals within Ru/SPS system. This dual action generates a critical photo-redox and crosslinking threshold that inhibits dityrosine formation until vitamin C is depleted. Thereby the threshold response needed for TVP is successfully established, which enables high-fidelity volumetric printing of native collagen. Post-printing construct densification (∼53% shrinkage) further improves feature resolution (80 µm positive; 120 µm negative) and yields mechanically stable and highly stretchable hydrogels (up to 180% strain). Collagen resin with vitamin C supports both cell seeding post-printing and cell-laden printing with high cell density and viability, enabling the rapid biofabrication of cell-instructive 3D microenvironments.

**Table of Contents (ToC):** 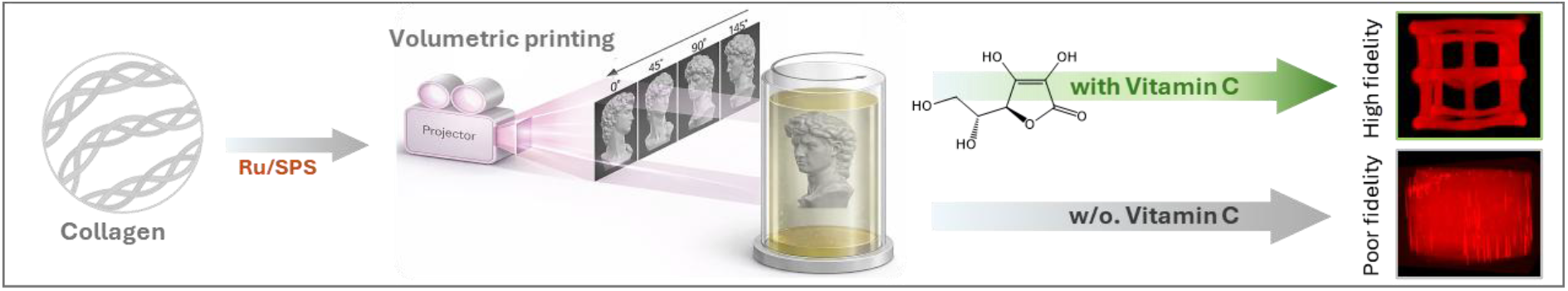

Tomographic volumetric printing (TVP) of native proteins is limited by uncontrolled background crosslinking. Here, vitamin C is introduced as a biocompatible redox-regulator to establish a tunable nonlinear polymerization threshold response for TVP. This strategy effectively suppresses background crosslinking and enables high-fidelity printing of pristine collagen. Subsequent post-print densification yields robust, elastic, and cell-compatible constructs with enhanced resolution for tissue engineering applications.

## 1. Introduction

The ability to fabricate biomaterials into complex, tissue-like 3D architectures is a central objective in tissue engineering and regenerative medicine.^[1,2]^ Among emerging biofabrication methods, Tomographic volumetric printing (TVP) has rapidly established itself as a transformative platform.^[3–5]^ Unlike traditional layer-by-layer approaches, TVP creates complex, centimeter-scale architectures in seconds by projecting optical tomograms into a rotating vial filled with light reactive resins. With its ultrafast speed, smooth surface and auxiliary support-free nature, TVP has been successfully applied to materials ranging from acrylates, epoxies, glass to hydrogel-based biomaterials.^[6–11]^ However, to date, TVP of hydrogels has predominantly relied on synthetic or chemically modified biomaterials, such as poly(ethylene glycol) diacrylate (PEGDA), methacrylated or norbornene-functionalized gelatin (GelMA, Gel-NB) and collagen (ColMA, Col-NB).^[7,9,12,13]^ In contrast, the direct printing of pristine proteins such as collagen, without chemical prefunctionalization, remains a challenge but is essential for preserving the intrinsic bioactivity of the native extracellular matrix.

One particularly attractive route toward printing unmodified protein-based biomaterials is to exploit their native tyrosine residues for crosslinking. Tyrosine residues, which are present in proteins such as collagen, fibrinogen, and silk, provide reactive phenolic groups that can be coupled to form dityrosine crosslinks.^[14–16]^ This crosslinking reaction can be induced *in vitro* using visible light (400–500 nm) via the ruthenium (II) tris-bipyridyl chloride (Ru(II)bpy_3_^2+^) and sodium persulfate (SPS) redox system (Ru/SPS).^[17,18]^ Because it eliminates the need for chemical modification, this approach simplifies bioink preparation and better preserves the biochemical features of the native tissue environment.

Despite its promise, translating dityrosine-based crosslinking to TVP presents a fundamental physicochemical limitation. A critical requirement for TVP is the presence of a desired non-linear threshold response, whereby the resin must remain liquid below a specific light exposure threshold but rapidly gel once that threshold is exceeded.^[11,19]^ Because light patterns must traverse the entire resin volume to build up the required dose in the target zone, non-target regions are repeatedly exposed to transmitted light. Without sufficient non-linear threshold behavior, background light accumulates and triggers crosslinking in unintended regions, severely compromising the accuracy and resolution of the printed structure. In free radical chain-growth photopolymerization systems, such as GelMA paired with a type I initiator^[2]^ like lithium phenyl 2,4,6 trimethyl (LAP), dissolved oxygen acts as a natural inhibitor, effectively providing this threshold response.^[20–22]^ While it is possible to engineer a limited, oxygen-dependent threshold in certain step-growth systems (e.g., thiol-ene networks combined with Type I initiators)^[7]^ by carefully tuning initiator concentrations to balance oxygen inhibition against initiation, this workaround is not applicable to dityrosine systems. Dityrosine crosslinking, when triggered by Ru/SPS (type II photoinitiator), relies on a step-growth mechanism that is largely oxygen-insensitive,^[18,23]^ leaving the system without an effective inhibitory mechanism. Specifically, under light illumination, SPS oxidizes Ru(II) to Ru(III), a potent oxidant, which near-immediate causes the formation of tyrosyl radicals and triggers premature crosslinking.^[18,24]^

Recent demonstrations of TVP using pristine biomaterials, such as pristine silk fibroin and silk sericin^[25]^ and decellularized extracellular matrix (dECM),^[26,27]^ have highlighted this challenge. These studies reported restricted irradiation windows and reduced geometric fidelity arising from insufficient suppression of background curing. While radical scavengers like TEMPO have been used as polymerization inhibitors to introduce nonlinear thresholding in other step-growth systems (e.g., thiol–ene chemistries)^[23,24]^, however, their poor aqueous solubility and cytotoxicity limit their compatibility with cell-laden bioinks.^[26]^

To overcome this limitation, we developed a redox-regulated approach utilizing vitamin C (L-ascorbic acid) to introduce the necessary threshold response into dityrosine crosslinking process. As a biocompatible sacrificial electron donor, vitamin C precisely modulates the Ru/SPS photo-initiating system by (i) reducing Ru(III) back to Ru(II) to sustain photocatalyst turnover, and (ii) scavenging persulfate radicals to suppress premature radical generation. This dual mechanism establishes a photo-redox threshold and thus effectively inhibits dityrosine crosslinking until vitamin C is locally depleted. This vitamin C regulated redox reaction process with a defined delay translates directly into the controlled, non-linear threshold response essential for the TVP. We designate this approach as the C-redox system.

To demonstrate the utility of the C-redox system for protein-based TVP, pristine collagen (type I) was selected as the model biomaterial. Collagen-I is the most abundant protein in mammals and a principal component of the extracellular matrix, which makes it an attractive bioink for engineering a broad range of tissue microenvironments.^[28]^ Under physiological conditions *in vitro*, collagen naturally self-assembles into native cross-banded fibrils (a process referred to as fibrillogenesis).^[29]^ This physical crosslinking process yields an opaque hydrogel, which is incompatible with optical printing techniques that require a liquid, transparent and low-scattering resin. To prevent fibril formation, collagen resin is conventionally maintained at low temperatures (e.g., 4 °C) or under acidic conditions.^[12]^ In this work, the collagen resin remained clear and printable by utilizing either an acidic aqueous solution (pH 3.5) or a neutral, glucose-rich buffer (pH 7.0) to inhibit fibril formation prior to printing.

Applied to TVP of native collagen, the C-redox system enables high-fidelity 3D printing by suppressing background curing, a level of control that is not achievable with Ru/SPS alone. This converts an intrinsically unstable redox chemistry into a practical and controllable printing strategy. We further show that the printed collagen constructs can be subjected to post-printing isotropic densification, which yields elastic hydrogels with a twofold increase in feature resolution. Taken together, the combination of precise patterning and post-printing densification underscores the versatility of the C-redox system for generating mechanically robust, biologically relevant protein constructs. In addition, the system supports direct cell encapsulation during bioprinting, broadening its potential for the fabrication of complex 3D tissue-like models and other engineered living materials.

## 2. Results

### 2.1. Regulation mechanism of vitamin C in the Ru/SPS redox system for dityrosine crosslinking inhibition

Pristine proteins, such as collagen, form dityrosine crosslinks when activated by the Ru/SPS photo-redox system (**Figure 1A**). To regulate this process, we introduced vitamin C, which undergoes oxidation upon electron donation (**Figure 1B**). We hypothesized that within the Ru/SPS system, vitamin C functions as a sacrificial electron donor to reduce photogenerated Ru(III) back to Ru(II) while scavenging persulfate radicals (**Figure 1C** and Figure S1, Supporting Information), and thereby inhibits dityrosine crosslinking.

**Figure 1.**
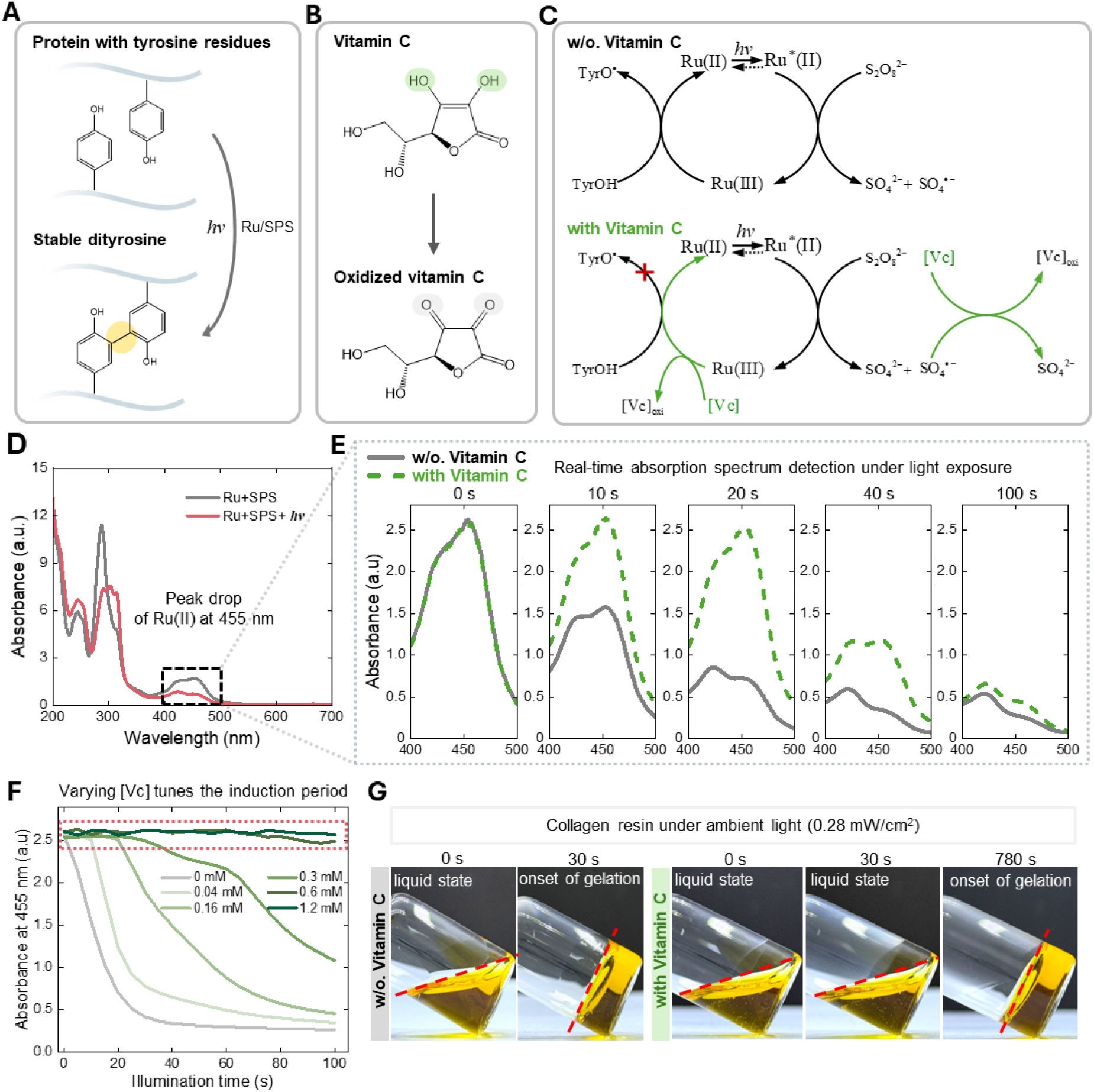
Regulation mechanism of vitamin C in the Ru/SPS redox system for inhibiting dityrosine crosslinking. **(A)** Schematic illustration of the formation of stable dityrosine crosslinks between protein chains mediated by the Ru/SPS photo-redox pair. **(B)** Chemical structures of vitamin C and its oxidized form. The green circles highlight the hydroxyl functional groups within the enediol group responsible for electron donation; oxidation results in the loss of electrons and protons. **(C)** Proposed reaction pathways for redox-induced tyrosine oxidation in the absence (top) and presence (bottom) of vitamin C. Vitamin C acts as a sacrificial electron donor, reducing Ru(III) back to Ru(II) and scavenging persulfate radicals. (**D)** UV-Vis absorption spectra of the Ru/SPS system before (gray line) and after (red line) light exposure, showing the characteristic bleaching of the MLCT peak at 455 nm (highlighted by the black box). **(E)** Real-time evolution of absorption spectra (400–500 nm) for Ru/SPS solutions without vitamin C (gray, solid) and with vitamin C (green, dashed) at indicated irradiation times (t=0, 10, 20, 40, 100 s). **(F)** Kinetics of Ru(II) absorbance decay monitored at 455 nm under continuous illumination with varying concentrations of vitamin C, demonstrating the tunable induction period marked by the red dot box. **(G)** Visual observation of the sol-gel transition in collagen resins containing Ru/SPS under ambient light (0.28 mW cm^-2^ at 405 nm). The addition of vitamin C significantly extends the liquid state duration compared to the control (without vitamin C).

To validate this proposed mechanism, we first established the baseline photochemical behavior of the Ru/SPS system without vitamin C. The aqueous solutions of Ru(II) exhibited a characteristic yellow color (Figure S2, Supporting Information), with distinct metal-to-ligand charge transfer (MLCT) absorption peaks at 455 nm and 285 nm.^[30]^ In contrast, aqueous SPS was colorless and showed no observable absorption in the visible range (300-700 nm). Upon mixing, the solution retained the spectroscopic signature of the Ru(II) complex. However, under light irradiation, the solution underwent photobleaching, accompanied by a decay of the absorption peaks at 455 nm and 285 nm (**Figure 1D**, and Figure S3, Supporting Information). This spectral change indicated the photo-oxidation of Ru(II) to Ru(III), a transition further supported by the emergence of a new absorption peak at 314 nm, characteristic of the oxidized species.^[31]^

To elucidate how vitamin C regulates this redox system, we performed kinetic absorption measurements under continuous light irradiation (2 mW cm^-2^ at 405 nm), using the 455 nm MLCT peak to track the Ru(II) concentration in real time. As shown in **Figure 1E**, in the absence of vitamin C, the 455 nm peak dropped significantly within the first 10 s and reaches a near-equilibrium state by 40 s, with no further changes observed by 100 s. However, the introduction of vitamin C fundamentally altered this kinetic profile. At t = 0 s, the absorption spectrum of the Ru/SPS/Vitamin C mixture overlaped with the Ru/SPS control, which confirmed that vitamin C (which absorbs primarily at 250 nm) did not interfere with the visible light absorption of the Ru(II) (**Figure 1E**, Figure S2, Supporting Information). Remarkably, under continuous irradiation, no significant drop in Ru(II) absorbance was observed at the 10 s or 20 s marks. This suggested that Ru(II) levels were maintained at a consistent steady-state, as the buildup of Ru(III) was effectively suppressed. It was only at the 40 s mark that the Ru(II) peak began to decline, which indicated that Ru(III) accumulates only after vitamin C has been sufficiently depleted (**Figure 1E**, Figure S3, Supporting Information). By 100 s, the absorbance of the vitamin C-regulated system converged with that of the control. These results confirmed that vitamin C acts as a sacrificial electron donor; it was rapidly consumed to reduce photogenerated Ru(III) back to the initial Ru(II) state, maintaining the Ru(III) concentration near zero while vitamin C was present.

Crucially, this chemically mediated delay before sustained oxidation established a distinct photo-redox threshold that gives rise to a non-linear kinetic response with a well-defined induction period. We demonstrated that the length of this induction period could be precisely tuned by varying the initial vitamin C concentration ([Vc]) (**Figure 1F**). Increasing the [Vc] linearly extended the duration of the Ru(II) plateau, which allowed for controllable timing of the oxidative onset. We compared this behavior to the well-known synthetic radical scavenger TEMPO, which also suppresses the oxidation of Ru(II) to Ru(III). On a molar basis, vitamin C yielded longer induction times than TEMPO (Figure S4, Supporting Information). This aligned with their respective redox stoichiometries, as vitamin C acts as a two-electron donor whereas TEMPO is a single-electron scavenger. Furthermore, beyond its molar efficiency, vitamin C also has the advantage of being a naturally occurring, non-toxic and highly water-soluble reagent (176 g/L at 20 °C) compared to TEMPO (9.7 g/L at 20 °C).

Notably, when the vitamin C concentration reached a stoichiometric equivalence with the SPS (2 mM in this study), the induction period became effectively infinite (Figure S5, Supporting Information). At this equivalence point, vitamin C fully suppressed sustained oxidation. Because vitamin C continuously regenerated photoactive Ru(II) from Ru(III), it indirectly drove the ongoing SPS consumption. At stoichiometric equivalence, both vitamin C and SPS were eventually exhausted, leaving no oxidant available to sustain the redox cycle. A phenomenon confirmed by the lack of peak decay across varying Ru concentrations (0.2 mM to 2 mM) (Figure S5, Supporting Information). This observation strongly supported the hypothesized dual regulatory role of vitamin C in the Ru(II)/SPS system. While it was experimentally established that vitamin C reduced Ru(III) back to Ru(II), the complete suppression of oxidation at stoichiometric equivalence strongly implied an additional radical-scavenging role that served as a secondary consumption pathway for vitamin C. Specifically, the depletion of SPS together with the absence of Ru(II) photobleaching indicated that vitamin C quenched both the oxidized metal center and the reactive sulfate radicals. This interpretation was consistent with the known antioxidant properties of vitamin C^[32]^ and was further supported by its superior efficacy compared to the synthetic, single-electron scavenger TEMPO (Figure S4, Supporting Information). Although direct radical scavenging was not independently measured here, the combined kinetic and stoichiometric data provided strong indirect evidence for this dual function.

Having established that vitamin C successfully suppressed the photo-redox reaction to create a tunable threshold, we next applied this C-redox system to tyrosine-rich (∼0.5 wt%) collagen^[33]^ resins to validate its ability to inhibit premature dityrosine crosslinking. We demonstrated this by monitoring collagen resins (pH=3.5, room temperature) exposed to ambient light (0.28 mW cm^-2^ at 405 nm, Table S1, Supporting Information). As expected, the control group (without vitamin C) reached the onset of gelation in just 30 s due to the lack of effective inhibitory mechanism. However, the addition of 0.6 mM vitamin C extended the stable liquid state to 780 s (**Figure 1G**). This represented a significant increase in the induction period before gelation. These findings showed that vitamin C introduces an inhibitory mechanism for dityrosine crosslinking by regulating the redox reaction.

### 2.2. C-redox creates tunable non-linear threshold response for high-fidelity TVP of collagen

TVP relies on the projection of optical patterns into a rotating resin to enable rapid generation of the intended 3D geometry through volumetric light-dose accumulation (**Figure 2A**). As established previously, the C-redox system created a tunable, chemically mediated photo-redox threshold (induction period) that functioned directly as a crosslinking threshold, which established the nonlinear threshold response required for TVP (**Figure 2B**).

**Figure 2.**
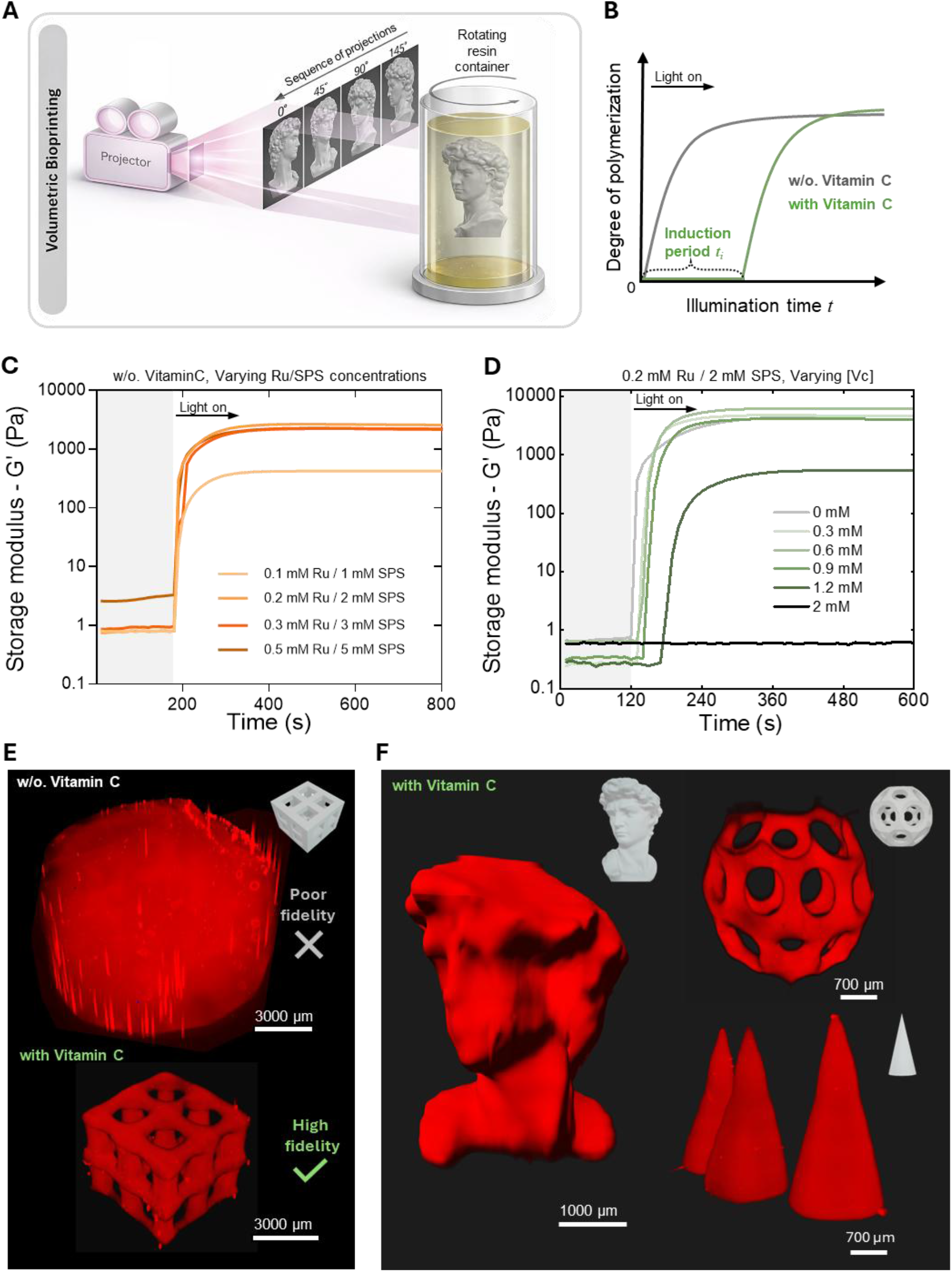
C-redox creates non-linear threshold response enabling high-fidelity volumetric printing of collagen. (**A)** Schematic of volumetric bioprinting: a dynamic light pattern (sequence of angular projections) is delivered by a laser source while the resin container rotates, enabling rapid 3D construction of the target geometry. (**B)** Non-linear threshold response curves. The addition of vitamin C (green) generates a distinct induction period (*t*_i_), whereas formulations lacking vitamin C (gray) exhibit an insufficient non-linear response. (**C)** Photoinitiator optimization for 5 mg/ml of collagen. Storage modulus G′ (Pa) vs time during illumination (“Light on”) for Ru/SPS (ruthenium/sodium persulfate) at 0.1/1, 0.2/2, 0.3/3, and 0.5/5 mM. 0.2/2 mM produced the highest plateau G′; higher Ru/SPS at the same ratio did not further increase the plateau. (**D)** Vitamin C titration at fixed photoinitiator concentrations. G′ vs time for formulations containing 0.2/2 mM Ru/SPS with 0–2 mM vitamin C. Vitamin C altered induction period *t*_i_ and gelation kinetics. (**E)** Representative print of a 2×2 lattice geometry printed with and without vitamin C. (**F)** High-fidelity volumetric prints produced with vitamin C, such as a model of David’s head, a C60 structure, and cones to demonstrate reproducibility and resolution.

Before this system was applied to TVP, photoinitiator (Ru/SPS) concentrations at a 1:10 ratio were tested by photorheology using a 405 nm light source.^[34]^ Across the tested Ru/SPS pairs (0.1/1, 0.2/2, 0.3/3, and 0.5/5 mM), storage modulus G′ increased sharply upon light exposure (“light on”), followed by a stable plateau (**Figure 2C**). The 0.1/1 mM Ru/SPS condition produced the lowest plateau modulus (plateau G′ ≈ 460 Pa). 0.2/2 mM Ru/SPS produced a plateau G′ of ≈ 2950 Pa, and further increases in Ru/SPS (0.3/3 and 0.5/5 mM) did not generate higher plateau moduli (Figure 2C). This suggested that the maximum storage modulus was limited by factors other than initiator availability (e.g., crosslinkable site mobility and density).^[35]^ Accordingly, 0.2/2 mM Ru/SPS was selected, and vitamin C was titrated (0–2 mM) to quantify its impact on gelation kinetics under otherwise identical illumination (**Figure 2D**). vitamin C produced a concentration-dependent delay in polymerization onset, corresponding to an increased induction time (*t*_i_) (**Figure 2D**). Between 0.3–0.9 mM, despite a progressively prolonged onset, the subsequent gelation rate was maintained at levels comparable to the control, allowing the gels to achieve high plateau moduli (G′ > 3 kPa). At higher vitamin C concentration (≥1.2 mM), gelation was substantially delayed but the plateau modulus decreased (G′ < 1 kPa), and at 2 mM gelation was completely suppressed over the experimental timescale (**Figure 2D**).

Building on this demonstration of a controlled induction period, we next sought to leverage this tunable crosslinking threshold response for volumetric printing of native collagen. To systematically evaluate how varying the vitamin C concentration affected print fidelity, we compared gear model prints formulated with 0, 0.3, 0.6, and 0.9 mM vitamin C (Figure S6, Supporting Information). While 0 and 0.3 mM were insufficient to prevent background curing, concentrations of 0.6 mM and 0.9 mM both successfully eliminated unintended crosslinking to yield highly resolved features. However, achieving this fidelity with 0.9 mM required a significantly higher light dose and prolonged print time (110 sec) compared to the 0.6 mM condition (65 sec) (Figure S6, Supporting Information). Consequently, 0.6 mM vitamin C was selected as the optimal formulation, offering an ideal balance between shape fidelity and fabrication speed.

Using this optimized formulation, we printed a 2×2 lattice model (**Figure 2E**). When printed without vitamin C, even at the lowest possible light dose of 68 mJ cm^?2^ (defined by the Tomolite TVP printer, Readily3D), the print showed pronounced background curing and a complete loss of feature definition. In contrast, with the addition of 0.6 mM vitamin C, lattices printed at approximately 600 mJ cm^−2^ showed well-defined struts and preserved overall architecture (**Figure 2E**). Furthermore, this optimized formulation reproducibly generated highly complex freeform geometries such as a model of David’s head, C60-like lattice, and conical structures within a printing time of 60-70 seconds (**Figure 2F** and Figure S7, Supporting Information). Collectively, these data show that vitamin C induced the desired nonlinear threshold response, to enable high-fidelity volumetric printing of pristine collagen.

To validate the pH-independence and generalizability of C-redox system, we performed photorheology experiments with collagen dissolved in phosphate-buffered saline (PBS) at physiological pH 7 (Supplementary Method 1, Supporting Information) and tested additional biomaterials, including fibrinogen pair with Ru/SPS and gelatin-based hydrogels (GelNB and GelSH) pair with LAP (Figure S8, Supporting Information). Vitamin C consistently modulated the induction period across the tested pH range (pH 3.5–7) and among all biomaterials examined, while nuclear magnetic resonance analysis confirmed its chemical stability in both acidic and neutral environments (Figure S9, Supporting Information).

### 2.3. Densification behavior and mechanical characterization of printed collagen constructs

Following TVP, collagen constructs (printed at acidic conditions) underwent a shrinkage process upon incubation in physiological conditions (PBS, 37 °C for 24 h). This “densification” was quantified, with printed cylinders for which both the lateral dimensions and the vertical axis were measured (**Figure 3A**). The hydrogels contracted to approximately 47% of their original linear size in both lateral and vertical directions (with no statistical difference between both direction), which corresponded to an isotropic linear shrinkage ratio of ∼ 53% (**Figure 3B** and Table S2, Supporting Information). This contraction could be attributed to the synergistic effects of osmotic pressure equilibration, pH-induced charge neutralization on collagen chains, and thermally driven polymer network reorganization.^[36]^

**Figure 3.**
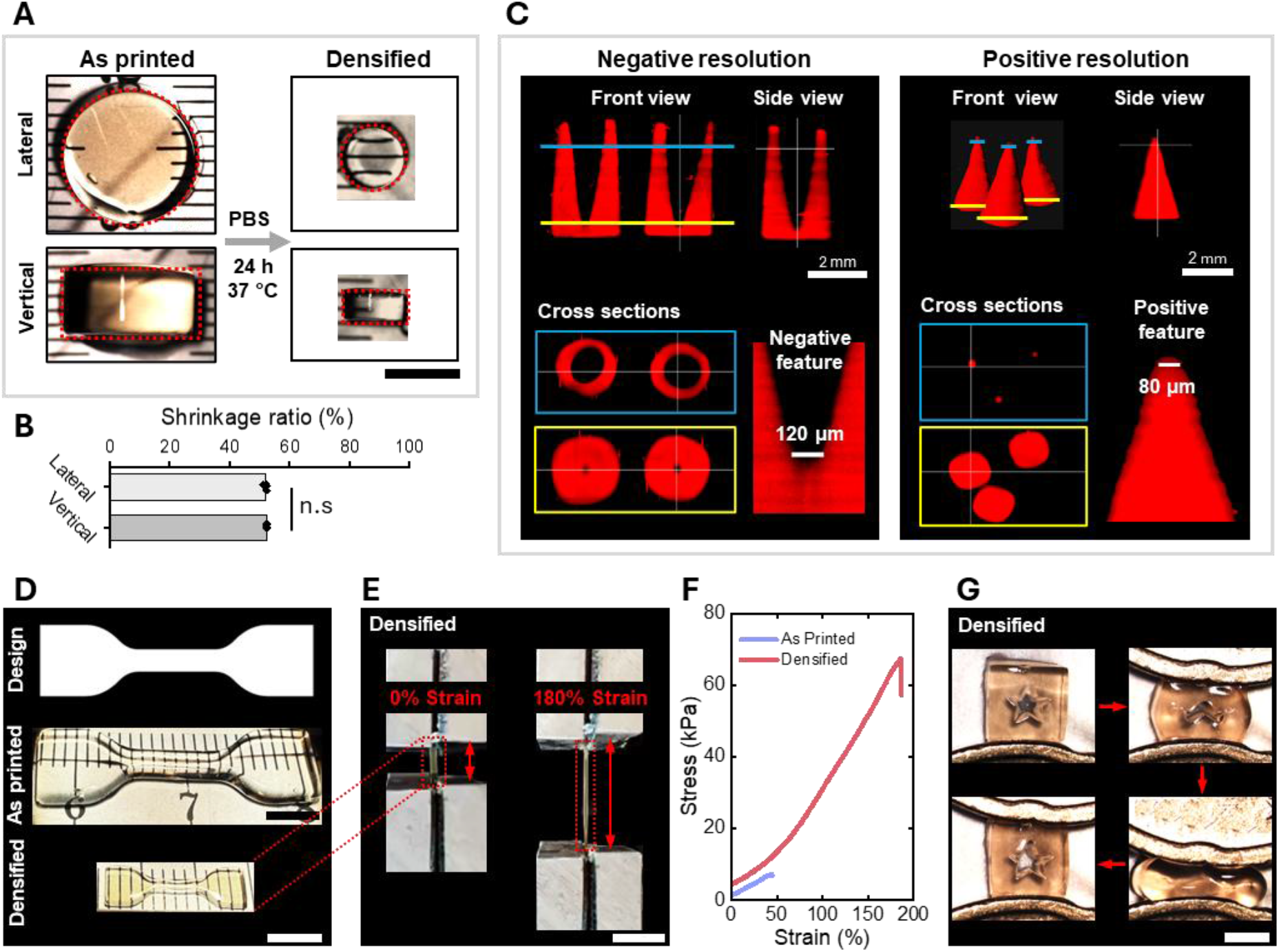
Densification behavior and mechanical characterization of printed collagen constructs. **(A)** Brightfield images of printed collagen cylinders before (“As printed”) and after incubation in PBS at 37 °C for 24 h (“Densified”). Top row: Lateral (top-down view); Bottom row: Vertical (side view). Red outlines indicate the gel boundaries. Scale bar: 5 mm. **(B)** Quantification of the shrinkage ratio in lateral and vertical dimensions. The shrinkage is isotropic (approx. 53%, n = 4; mean ± SD) with no significant difference (n.s.) between the Lateral and Vertical directions. **(C)** Resolution analysis of densified structures utilizing the shrinkage to enhance detail. Left: Negative resolution test showing open void channels with a minimum feature size of 120 µm. Right: Positive resolution test showing solid cone tips with a minimum feature size of 80 µm. **(D)** Geometric fidelity across scales. Top: Digital CAD design of a dogbone specimen. Middle: As-printed construct. Bottom: Densified construct, illustrating uniform, distortion-free shrinkage. Scale bars: 5 mm. **(E)** Tensile testing snapshots of a densified dogbone specimen at 0% strain (relaxed) and 180% strain (stretched), demonstrating high extensibility. Scale bar: 5 mm. **(F)** Comparison of tensile stress-strain curves of dogbone specimens between As-printed (blue curve) and Densified sample (red curve). **(G)** Cyclic compression test of a densified hydrogel containing a star-shaped internal void. The sample withstands 60% compression and instantaneously recovers its original shape without structural damage. Scale bar: 2 mm.

We leveraged this isotropic densification as post-print resolution enhancement strategy to achieve fine feature sizes. To quantify the resolution limits, we printed cone structures to assess both positive (solid) and negative (void) features. Following densification, positive features were as fine as 80 µm and negative features (inverted cones) were resolved down to 120 µm. (**Figure 3C**). To evaluate the implications of densification on mechanical performance, dogbone structures were printed for tensile testing and underwent the same isotropic shrinkage as the disks (**Figure 3D**). Tensile testing revealed that densification altered the failure behavior of the hydrogel. While as-printed samples failed at approximately 50% strain, densified samples exhibited greatly improved extensibility; withstood strains of up to180% (**Figure 3E**) and enduring significantly higher stress compared to the control without failure (**Figure 3F**, Movie S1-3, Supporting Information). The densified constructs possessed a tensile modulus of approximately 15 kPa (n = 5, extracted from the 5-20% strain region, Figure S10, Supporting Information). This increase in stretchability and load-bearing capacity suggested that the densification process effectively promoted network compaction and physical entanglement.^[34]^

Finally, the compressive resilience of a densified collagen construct containing a complex star-shaped internal void was assessed. The structure withstood 60% compression and rapidly recovered its original geometry and the printed internal void, upon release (**Figure 3G**). Together with its high fidelity, extensibility and elasticity, this underscored the potential of densified collagen for mechanically demanding tissue engineering applications.

### 2.4. TVP-printed collagen supports structural retention, post-seeding, and cell-laden printing of 3T3 fibroblast

Next, it was evaluated whether TVP-printed collagen constructs could provide a stable and cytocompatible microenvironment for cells by post-printing seeding with fibroblasts. Across a 7-day culture period, printed disks retained their macroscopic geometry (**Figure 4A**). Quantification of matrix integrity, assessed by disk diameter of cell-seeded constructs relative to acellular controls, showed no change over time (n = 3, mean ± SD; **Figure 4B**), which indicated limited bulk degradation and no cell-driven contraction within this timeframe.

**Figure 4.**
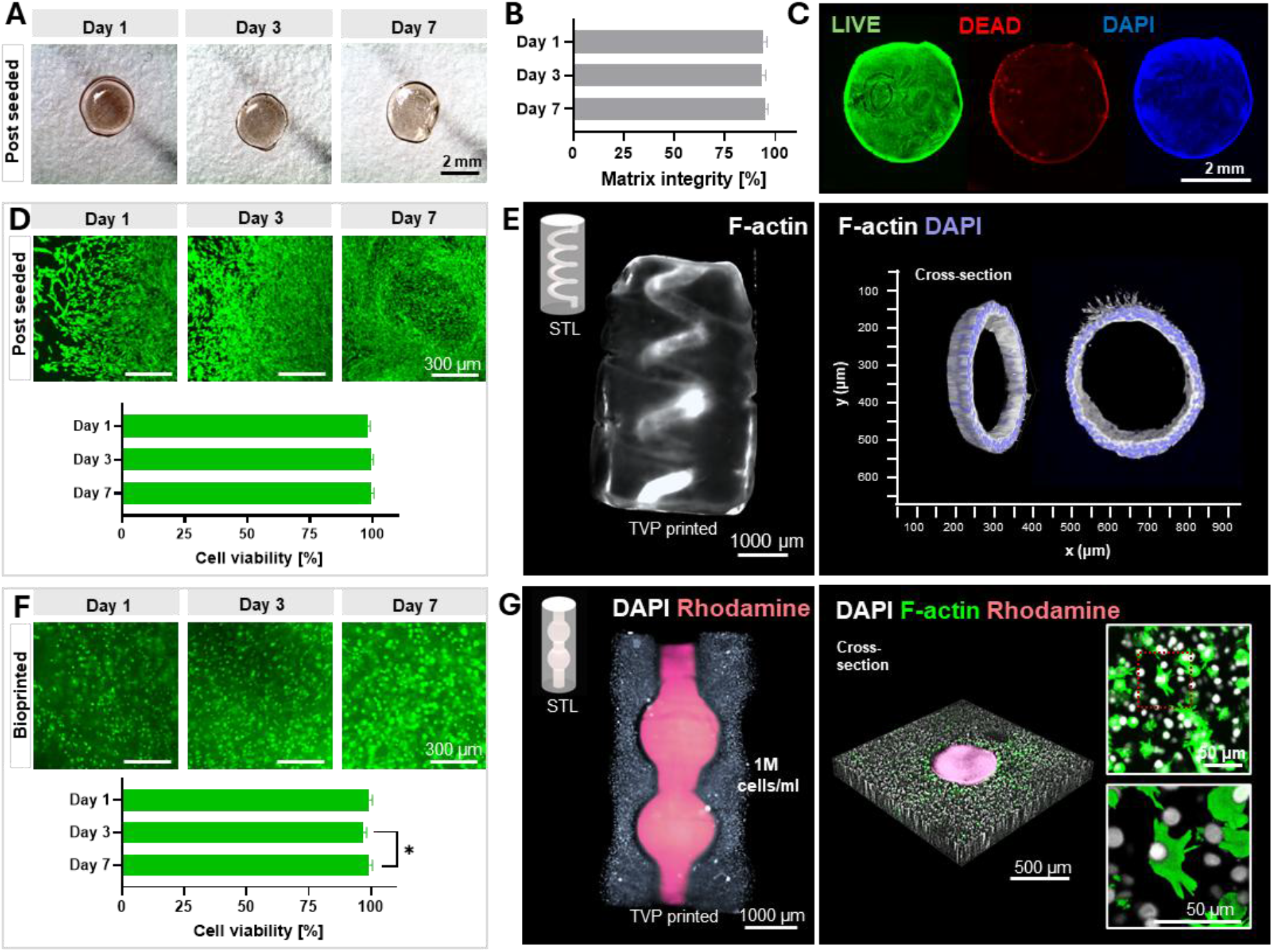
Cytocompatibility and structural retention of TVP-printed collagen constructs with post-seeded and bioprinted 3T3 fibroblasts. (**A)** Brightfield images of printed collagen disks at Day 1, 3, and 7 showing preserved geometry. (**B)** Quantification of construct stability/degradation (matrix integrity) expressed as projected disk diameter over time (n = 3; mean ± SD). (**C)** Corresponding whole-disk viability imaging at Day 7 confirms that constructs maintain macroscopic integrity and also support cell viability (LIVE: calcein AM, green; DEAD: propidium iodide, red; nuclei: DAPI, blue). (**D)** Representative calcein AM fluorescence images of post-seeded 3T3 fibroblasts on printed collagen disks at Day 1, Day 3, and Day 7, with corresponding quantification (n = 3; mean ± SD). (**E)** 3D reconstruction of F-actin organization in the printed helical collagen construct after post-seeding via injection of 3T3 fibroblasts into the helical channel (left). Corresponding cross-sectional F-actin (red) and DAPI (blue) images (right) demonstrate cellular adhesion and distribution along the collagen wall. (**F)** Representative calcein AM fluorescence images of bioprinted (cell-laden) collagen disks at Day 1, Day 3, and Day 7, with corresponding quantification (n = 3; mean ± SD; *p < 0.05). (**G)** Printed ductal–alveolar model with a collagen stromal compartment containing 3T3s (1 × 10^6^ cells mL^−1^; DAPI, white); the central lumen was perfused with rhodamine-labeled GelMA (pink) to visualize channel architecture, with insets highlighting F-actin organization and nuclear distribution after 1 day.

Because matrix stability is only biologically meaningful if the cells remain viable and able to interact with the construct, cell viability and attachment were also assessed. Whole-disk live/dead staining confirmed rapid 3T3 attachment (within 24 h) and progressive spreading across the disk surface over 7 days, with strong calcein AM signal and minimal propidium iodide staining throughout the culture period on collagen printed in both acidic and neutral conditions in glucose rich buffer (**Figure 4C–D**, Figure S11A-B, Supporting Information). Brightfield top- and side-view imaging confirmed structural integrity in both printing conditions after 1 week of culture relative to acellular controls (Figure S11D–E, Supporting Information). To assess cell seeding and culture within geometrically complex collagen architectures, a helix within a cylinder was printed and fibroblasts were post-seeded by injection into the helical channel (**Figure 4E**). After seven days of culture, F-actin staining showed that cells occupied the helical lumen while the printed geometry was preserved. Additional high-magnification cross-sectional channel views verified robust cell adhesion and lining of the lumen perimeter with F-Actin expressing fibroblasts (**Figure 4E**).

In addition to post-seeding, cell-laden bioprinting directly incorporated living cells into printed 3D matrices. To assess the suitability of our collagen resin and printing process with cell-laden printing, fibroblasts were resuspended in the collagen resin and printed as disk constructs, that were washed in PBS and cell culture medium immediately after printing. Over a culture period of 7 days, high viability was maintained, as shown by live/dead imaging and quantification at days 1, 3, and 7 (**Figure 4F**, Figure S11C, Supporting Information).

Next, fibroblasts were resuspended in the collagen resin at 1 × 10^6^ cells ml^−1^ to assess whether densely cellular stromal compartments could be fabricated while preserving shape fidelity. A ductal-alveolar channel was printed, and the hollow structures were perfused with rhodamine-labeled GelMA to visualize the shape. Rhodamine signal delineated the intended lumen geometry, while DAPI staining (white) showed nuclei (cells) homogenously distributed throughout the printed collagen matrix. In addition, cells displayed F-actin signal (green) and early spreading after 24 h, consistent with early cytoskeletal remodeling and matrix engagement, within the printed collagen network (**Figure 4G**, right).

## 3. Discussion and Conclusion

In this study, we demonstrated that vitamin C introduces an inhibitory mechanism for dityrosine crosslinking by regulating Ru/SPS redox reactions. As a naturally occurring, biocompatible, and water-soluble redox regulator, vitamin C suppressed Ru(III) accumulation and scavenged persulfate radicals to establish a photo-redox threshold. In protein-based resins, the developed C-redox system created a chemical mediated crosslinking threshold that functioned directly as the non-linear threshold response required for volumetric printing. Ultimately, these findings enabled the high-fidelity TVP of pristine collagen.

After printing, collagen densification at 37°C in PBS was leveraged as a post-printing resolution enhancement strategy, for improved feature definition (resulting positive feature: 80 µm, negative feature: 120 µm). Similar approaches have been previously reported, where an initially water-rich network was compacted to sharpen features and increased effective polymer density.^[37]^ In the context of TVP, this shrinkage-based refinement is particularly advantageous. Because TVP relies on coordinated multi-angle illumination to build dose contrast between the target object and the background, achieving ultra-fine resolution can be optically challenging due to factors such as light scattering and attenuation. While recent computational and optical advances, such as projection optimization and dose extraction via secondary light path have made significant progress in addressing this,^[38–42]^ our densification process serves as a highly effective, complementary material-driven strategy. By bridging these advanced optical techniques with post-print shrinkage, the feature resolution and fidelity of volumetrically printed protein-based hydrogels can be further maximized.^[38–42]^ Moreover, densification significantly enhanced the mechanical performance of collagen hydrogels, which rendered them highly elastic and capable of withstanding strains up to 180% without failure. This positions vitamin C printed densified collagen scaffolds ideal candidates for tissue engineering applications, particularly in mechanically demanding tissues like muscles and tendons, as well as for applications in soft robotics. Densification introduces a predictable but substantial size change that may require pre-compensation in the CAD/printing energy-map when the final part dimensions must match a specified target (e.g., channel diameters, wall thicknesses, or connector interfaces). Here, the extent and rate of shrinkage are expected to depend on formulation and processing variables (buffer composition, ionic strength/osmolarity, pH, and temperature/time) so these parameters require tight control to ensure reproducible dimensions across different geometries and length scales.

Beyond print fidelity and mechanical performance, our results showed that C-redox printed collagen supports both post-seeding and cell-laden biofabrication, highlighting the broader biological utility of this platform. TVP-printed collagen constructs fabricated from acidic precursor solutions supported cell attachment, and remained structurally stable over one week in culture. At the same time, future translation will require careful optimization of the collagen solution, as the physical crosslinking and cell behavior are strongly influenced by pH, temperature, ionic strength, collagen concentration and collagen I type (i.e., telopeptide-containing collagen versus atelocollagen).^[43]^ Printing parameters, including initiator concentration, vitamin C concentration, and light dose, will likely require cell-type-specific tuning, while collagen content should be adjusted to better match the density and mechanics of target tissues. Longer-term studies should also address cell-driven remodeling and mass-transport limitations in larger or more densely populated constructs.^[44]^

Although this study focused on collagen, the underlying inhibitory mechanism introduced by C-redox may be transferable to other step-growth-oriented biomaterials. This possibility remains to be demonstrated experimentally, but it raises the prospect of extending thresholded crosslinking to a broader range of protein-based and modified biomaterial systems like dECM and fibrinogen, as well as pre-functionalized proteins and tyramine-modified polymers such as Gel-SH, Gel-NB, and tyramine-modified hyaluronic acid (HA-Tyr). Such validation would position the C-redox system as a versatile platform for controlled photopolymerization across photolithographic systems such as TVP, Holographic printing,^[42–45]^ and (bio)Xolography,^[46]^ where controlled crosslinking and inhibition of unintended reactions are critical.

## 4. Experimental Section/Methods

### UV-Vis Spectrophotometry

The absorption spectra of aqueous solutions containing Ru(II), SPS, vitamin C, and their various combinations were measured using a NanoDrop spectrophotometer (Thermo Scientific). Measurements were conducted under two distinct operational modes. For standard spectral characterization, a 2 μL sample was loaded to measure absorbance across a wavelength range of 200–700 nm. For *in situ* kinetic measurements under light exposure, 1.2 mL of the sample was loaded into a 10 mm pathlength cuvette. Illumination was provided by a custom-built platform equipped with a 405 nm LED (M405L4, Thorlabs) and a collimation lens with a 100 mm focal length (AC254-100-AB, Thorlabs). The light intensity was set to 2 mW cm^-2^. Kinetic data was recorded at 5-second intervals during continuous illumination, capturing an effective UV-Vis range of 300–700 nm.

### Preparation of collagen photoresin

Medical-grade purified type I atelocollagen powder of bovine origin (CBPE2, Symatese) was dissolved in ultrapure water (ELGA LabWater) to a final concentration of 5 mg/mL. The mixture was incubated at 4 °C for 24 hours, yielding a clear aqueous solution with a pH of approximately 3.5. To complete the photoresin, a photoinitiator system comprising 0.2 mM ruthenium (Ru) and 2 mM sodium persulfate (SPS) (Advanced BioMatrix) was added to the solution. For formulations incorporating vitamin C (95210, Sigma-Aldrich), 0.6 mM vitamin C (different concentration as indicated) was added to the collagen resin prior to the introduction of the Ru/SPS system. To remove bubbles formed during resin mixing, the mixture was centrifuged at 500 rpm and 20 °C for 5 minutes.

### Photorheology

Rheology was performed using an Anton Paar MCR 302e rheometer with a 20 mm parallel plate geometry and a glass floor. The rheometer was combined with the Omnicure Series1000 lamp (Lumen Dynamics) with sequential 400–500 nm and narrow 405 nm bandpass filters (Thorlabs). Measurements were performed in the dark for 2 or 3 min before irradiating the sample with 405 nm light at 4 mW cm^−2^ irradiance. 76 µL of the sample was loaded onto the rheometer, with a gap distance of 0.2 mm. Modulus was recorded at 10-second intervals during the measurements.

### Volumetric Printing

0.9 mL of the collagen formulations were loaded into 10 mm printing vials at room temperature. Tomographic projections were performed on a commercially available printer in the dark with voxel resolution of 25 µm (Tomolite, Readily 3D SA).

### Print Post-processing and Densification

Following fabrication, the printed constructs were immersed in distilled water to extract unreacted resin. To ensure thorough washing, the water was replaced three times over a 2-hour period. The prints were subsequently transferred into a phosphate-buffered saline (PBS) solution and incubated at 37 °C for 24 hours to induce densification. For samples designated for light-sheet microscopy imaging, a fluorescent staining step was introduced after the initial wash but prior to densification. Specifically, 0.1% (v/v) methacryloxyethyl thiocarbamoyl rhodamine B was added to the water bath. The prints were left to incubate in this solution overnight, allowing the rhodamine B to diffuse into the network and impart a transparent red color, after which the standard thermal densification process in PBS was carried out.

### Print Shrinkage Ratio Characterization

The shrinkage ratio was quantified separately for the lateral and vertical directions using the following formula:

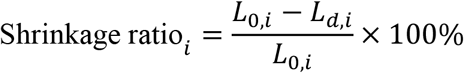

Where: *L*_0,*i*_ is the initial dimension in the (i)-th direction (lateral or vertical), *L*_*d,i*_ is the corresponding dimension after densification, and *i* denotes either the lateral or vertical plane.

### Tensile Testing

Dogbone specimens for tensile testing were designed based on the ISO 527-2 standard, with dimensions adapted to accommodate the projection plane of the Tomolite printer. The specimens were fabricated within a glass cuvette (2 mm thickness) using single-image projection at a light dose of 430 mJ/cm^2^. Uniaxial tensile tests were performed using a Texture Analyzer (TA.XTplus, Stable Micro Systems). To prevent slippage and ensure the hydrogel was securely held, the clamping device was lined with sandpaper. Testing was conducted at a constant displacement rate of 0.1 mm/s up to a target strain of 300%. Stress-strain curves were generated from the recorded data, from which the tensile stiffness was extracted and reported.

### Light sheet microscopy

An axially scanned light sheet microscope (MesoSPIM, V4) was used to image printed constructs. The constructs were imaged in a square glass cuvette with PBS which was inserted into a custom 3D-printed sample holder and submerged in a quartz chamber filled with miliQ water, mounted onto the MesoSPIM microscope stand. For imaging, a macro-zoom system (Olympus MVX-10) and 2x or 5x air objective (Olympus MVPLAPO1x) were used. Voltage adjustments using the electrically tunable lens (ETL) were performed. Step sizes were 5 or 10 µm.

### Cell culture

NIH 3T3 murine fibroblasts (ATCC) were cultured in DMEM with GlutaMAX medium supplemented with 10% Fetal Bovine Serum (FBS) and 5ml of Pen/Strep. Cells were expanded in tissue culture flasks at 37 °C and 5% CO_2_. The cells were passaged at 80% confluency using 0.25% w/v trypsin.

### Scaffold post-seeding

For post-print seeding, 3T3 fibroblasts were prepared at 1 × 10^6^ cells mL^−1^ and injected into the printed helical channel using a 20 µL pipette. Seeded scaffolds were transferred to a well plate and incubated for 45 min to allow cell attachment before gently adding culture medium to fill the well.

### Cell-laden volumetric bioprinting

For cell-laden printing, 1 × 10^6^ 3T3 fibroblasts were pelleted and resuspended in 1 mL collagen to generate a homogeneous cell suspension. Photoinitiator (Ru/SPS) and vitamin C were added immediately before printing. Cell-laden constructs were volumetrically printed, rinsed with cold PBS, transferred to culture medium, and maintained in a humidified incubator for subsequent culture. Ducatl-alveolar channels were subsequently perfused with fluorescently labeled GelMA and cured in a UV-box (BSL-01, Opsytec Dr. Gröbel GmbH).

### Matrix integrity and viability

For post-seeding, 2 × 10^4^ cells in 10 µL medium were pipetted onto each VP-printed and postprocessed disk and allowed to adhere for 45 min (37 °C) before gently adding culture medium to fill the well. Disk diameter was measured at days 1, 3, and 7 (n = 3 disks; 3 measurements per disk) and normalized to acellular controls to quantify matrix integrity. At the same time points, viability was assessed using calcein AM (live), propidium iodide (PI; dead), and DAPI (nuclei). Viability was quantified as PI^+^/DAPI^+^ cells and reported as percent. Cell-laden constructs were analyzed using the same workflow.

### Immunohistochemistry and confocal imaging

Constructs with cells were fixed in 4% PFA for 2h, and then washed 3 times in PBS, and blocked with 5% bovine serum albumin (BSA, Millipore Sigma) with 0.2% Triton-X100 (Sigma, T8787-100ml) in PBS for 35 min at room temperature. The constructs were incubated with Hoechst and F-Actin (abcam, ab176753 & ab176756) for 3 hours at RT washed 3 times in PBS and imaged using a Leica TCS SP8 confocal microscope.

### Statistical analysis

Quantification and statistical analysis. All experiments were performed with at least three replicates, unless otherwise specified. The sample size (n) for each experiment is indicated in the corresponding figure legends. Data analysis was performed using Microsoft Excel (Microsoft 365 MSO, Version 2412, Build 16.0.18324.20092, 64-bit), MATLAB R2018a (9.4.0.813654), GraphPad Prism (v10.4.0, Windows), OriginPro2024 and Fiji/ImageJ 1.54f (Java 1.8.0_332). Unless otherwise stated, data are presented as mean ± SD, and statistical significance was assessed using a paired t-test. Differences were considered statistically significant at P < 0.05.

### Statement of AI usage

ChatGPT (OpenAI) and Gemini (Google) were used to refine the language of this manuscript, including spelling and grammar. Figure 2A was enhanced using AI based on our original design. No AI tools were used to generate scientific research results.

## Conflict of Interest Statement

B.W., A.H., D.F., and M.Z.W. are inventors on a PCT patent application related to this work (PCT/EP26171225). All authors declare that they have no other competing interests.

## Acknowledgements

M.Z.-W. gratefully acknowledges funding from Swiss National Science Foundation, Bridge Discovery (40B2-0_211764) and the ETH Foundation (23-1 ETH-12). We thank Annalena Maier, Isabel Hui, Michael Winkelbauer and all the members of the Tissue Engineering and Biofabrication laboratory for the helpful discussions and inputs. The authors gratefully acknowledge ScopeM for their support in this work. Lightsheet imaging was performed with equipment maintained by the Center for Microscopy and Image Analysis, University of Zurich.

## Data Availability Statement

All data needed to evaluate the conclusions in the paper are present in the paper and/or the Supplementary Materials. Data also is available in the ETH Zurich Research Collection (https://doi.org/10.3929/ethz-c-000797572) under the terms of the repository’s data-sharing policies.

## Notes

**Conflict of Interest:** B.W., A.H., D.F., and M.Z.W. are inventors on a PCT patent application related to this work (PCT/EP26171225). All authors declare that they have no other competing interests.

https://doi.org/10.3929/ethz-c-000797572

## References

[1] C.-F. He, T.-H. Qiao, G.-H. Wang, Y. Sun, Y. He, Nat Rev Bioeng 2025, 3, 143.

[2] P. Chansoria, R. Rizzo, D. Rütsche, H. Liu, P. Delrot, M. Zenobi-Wong, Chem. Rev. 2024, 124, 8787.

[3] B. E. Kelly, I. Bhattacharya, H. Heidari, M. Shusteff, C. M. Spadaccini, H. K. Taylor, Science 2019, 363, 1075.

[4] D. Loterie, P. Delrot, C. Moser, Nat Commun 2020, 11, 852.

[5] P. N. Bernal, S. Florczak, S. Inacker, X. Kuang, J. Madrid-Wolff, M. Regehly, S. Hecht, Y. S. Zhang, C. Moser, R. Levato, Nat Rev Mater 2025, 10, 826.

[6] B. Wang, E. Engay, P. R. Stubbe, S. Z. Moghaddam, E. Thormann, K. Almdal, A. Islam, Y. Yang, Nat Commun 2022, 13, 367.

[7] R. Rizzo, D. Ruetsche, H. Liu, M. Zenobi-Wong, Advanced Materials 2021, 33, 2102900.

[8] Q. Thijssen, A. Quaak, J. Toombs, E. De Vlieghere, L. Parmentier, H. Taylor, S. Van Vlierberghe, Advanced Materials 2023, 35, 2210136.

[9] P. N. Bernal, M. Bouwmeester, J. Madrid-Wolff, M. Falandt, S. Florczak, N. G. Rodriguez, Y. Li, G. Größbacher, R.-A. Samsom, M. van Wolferen, L. J. W. van der Laan, P. Delrot, D. Loterie, J. Malda, C. Moser, B. Spee, R. Levato, Adv Mater 2022, 34, e2110054.

[10] P. N. Bernal, P. Delrot, D. Loterie, Y. Li, J. Malda, C. Moser, R. Levato, Advanced Materials 2019, 31, 1904209.

[11] J. Madrid-Wolff, J. Toombs, R. Rizzo, P. N. Bernal, D. Porcincula, R. Walton, B. Wang, F. Kotz-Helmer, Y. Yang, D. Kaplan, Y. S. Zhang, M. Zenobi-Wong, R. R. McLeod, B. Rapp, J. Schwartz, M. Shusteff, H. Talyor, R. Levato, C. Moser, MRS Commun 2023, 13, 764.

[12] M. Winkelbauer, A. Hasenauer, D. Rütsche, H. Liu, J. Janiak, M. Nguyen, K. L. Christman, M. Zenobi-Wong, P. Chansoria, Advanced Healthcare Materials 2025, 14, 2405105.

[13] D. Ribezzi, J.-P. Zegwaart, T. Van Gansbeke, A. Tejo-Otero, S. Florczak, J. Aerts, P. Delrot, A. Hierholzer, M. Fussenegger, J. Malda, J. Olijve, R. Levato, Advanced Materials 2025, 37, 2409355.

[14] B. P. Partlow, M. B. Applegate, F. G. Omenetto, D. L. Kaplan, ACS Biomater Sci Eng 2016, 2, 2108.

[15] M. B. Maina, Y. K. Al-Hilaly, L. C. Serpell, Front. Neurosci. 2023, 17, DOI 10.3389/fnins.2023.1132670.

[16] M. Correia, M. T. Neves-Petersen, P. B. Jeppesen, S. Gregersen, S. B. Petersen, PLOS ONE 2012, 7, e50733.

[17] A. Gulzar, E. Yildiz, H. N. Kaleli, M. A. Nazeer, N. Zibandeh, A. N. Malik, A. Y. Taş, I. Lazoğlu, A. Şahin, S. Kizilel, Acta Biomater 2022, 147, 198.

[18] D. A. Fancy, T. Kodadek, Proceedings of the National Academy of Sciences 1999, 96, 6020.

[19] B. Wang, W. S. H. Safari, S. Kaveh Hedayati, J. P. C. Narag, T. D. V. Christiansen, A. A. Schiefler, H. O. Sørensen, J. R. Frisvad, K. Almdal, A. Islam, Y. Yang, 2023, DOI 10.48550/arXiv.2303.13941.

[20] S. C. Ligon, B. Husár, H. Wutzel, R. Holman, R. Liska, Chem Rev 2014, 114, 557.

[21] C. Decker, A. D. Jenkins, Macromolecules 1985, 18, 1241.

[22] Y. Zhang, K. Houlahan, D. Webber, N. Milliken, K. L. Sampson, H. W. de Haan, H. Li, R. Vlaming, L. Gaburici, A. Orth, C. Paquet, Advanced Materials 2025, 37, e08729.

[23] C. C. Cook, E. J. Fong, J. J. Schwartz, D. H. Porcincula, A. C. Kaczmarek, J. S. Oakdale, B. D. Moran, K. M. Champley, C. M. Rackson, A. Muralidharan, R. R. McLeod, M. Shusteff, Adv Mater 2020, 32, e2003376.

[24] P. J. Wright, A. M. English, J Am Chem Soc 2003, 125, 8655.

[25] M. Xie, L. Lian, X. Mu, Z. Luo, C. E. Garciamendez-Mijares, Z. Zhang, A. López, J. Manríquez, X. Kuang, J. Wu, J. K. Sahoo, F. Z. González, G. Li, G. Tang, S. Maharjan, J. Guo, D. L. Kaplan, Y. S. Zhang, Nat Commun 2023, 14, 210.

[26] A. Hasenauer, K. Bevc, M. C. McCabe, P. Chansoria, A. J. Saviola, K. C. Hansen, K. L. Christman, M. Zenobi-Wong, Sci Adv 2025, 11, eadu5793.

[27] L. Lian, M. Xie, Z. Luo, Z. Zhang, S. Maharjan, X. Mu, C. E. Garciamendez-Mijares, X. Kuang, J. K. Sahoo, G. Tang, G. Li, D. Wang, J. Guo, F. Z. González, V. Abril Manjarrez Rivera, L. Cai, X. Mei, D. L. Kaplan, Y. S. Zhang, Advanced Materials 2024, 36, 2304846.

[28] Z. Zhang, Z. Ren, S. Chen, X. Guo, F. Liu, L. Guo, N. Mei, Arch Toxicol 2018, 92, 717.

[29] K. Tarnutzer, D. Siva Sankar, J. Dengjel, C. Y. Ewald, Sci Rep 2023, 13, 4490.

[30] D. M. Darvish, Mater Today Bio 2022, 15, 100322.

[31] C. Daul, E. J. Baerends, P. Vernooijs, Inorg. Chem. 1994, 33, 3538.

[32] B. Limburg, E. Bouwman, S. Bonnet, ACS Catal. 2016, 6, 5273.

[33] S. J. Padayatty, A. Katz, Y. Wang, P. Eck, O. Kwon, J.-H. Lee, S. Chen, C. Corpe, A. Dutta, S. K. Dutta, M. Levine, J Am Coll Nutr 2003, 22, 18.

[34] K. S. Lim, B. S. Schon, N. V. Mekhileri, G. C. J. Brown, C. M. Chia, S. Prabakar, G. J. Hooper, T. B. F. Woodfield, ACS Biomater. Sci. Eng. 2016, 2, 1752.

[35] A. Nunes, A. Pramanick, D. Kelly, R. Ramakrishnan, V. Sergis, V. K. Doan, H. A. Tran, K. S. Lim, H. Almeida, A. Daly, 2026.

[36] D. Iudin, L. J. J. A. Gerridzen, P. N. Bernal, C. C. L. Schuurmans, M. Neumann, L. Nguyen, M. J. van Steenbergen, J. Hak, W. Li, C. Casadidio, A. M. van Genderen, R. Masereeuw, R. Levato, Y. S. Zhang, B. G. P. van Ravensteijn, T. Vermonden, Biomacromolecules 2025, 26, 4108.

[37] J. Gong, C. C. L. Schuurmans, A. M. van Genderen, X. Cao, W. Li, F. Cheng, J. J. He, A. López, V. Huerta, J. Manríquez, R. Li, H. Li, C. Delavaux, S. Sebastian, P. E. Capendale, H. Wang, J. Xie, M. Yu, R. Masereeuw, T. Vermonden, Y. S. Zhang, Nat Commun 2020, 11, 1267.

[38] A. Orth, D. Webber, Y. Zhang, K. L. Sampson, H. W. de Haan, T. Lacelle, R. Lam, D. Solis, S. Dayanandan, T. Waddell, T. Lewis, H. K. Taylor, J. Boisvert, C. Paquet, Nat Commun 2023, 14, 4412.

[39] S. Florczak, G. Größbacher, D. Ribezzi, A. Longoni, M. Gueye, E. Grandidier, J. Malda, R. Levato, Nature 2025, 645, 108.

[40] D. Webber, Y. Zhang, M. Picard, J. Boisvert, C. Paquet, A. Orth, Opt. Express, OE 2023, 31, 5531.

[41] J. Madrid-Wolff, A. Boniface, D. Loterie, P. Delrot, C. Moser, Advanced Science 2022, 9, 2105144.

[42] M. I. Álvarez-Castaño, A. G. Madsen, J. Madrid-Wolff, V. Sgarminato, A. Boniface, J. Glückstad, C. Moser, Nat Commun 2025, 16, 1551.

[43] E. O. Osidak, V. I. Kozhukhov, M. S. Osidak, S. P. Domogatsky, Int J Bioprint 2020, 6, 270.

[44] M. Pires Figueiredo, S. Rodríguez-Fernández, F. Copes, D. Mantovani, NPJ Biomed Innov 2025, 2, 16.

[45] X. Wang, Y. Ma, Y. Niu, B. Xiong, A. Zhang, G. Zhang, Y. Chen, W. Wei, L. Fang, J. Wu, Q. Dai, Nature 2026, 650, 882.

[46] A. Wolfel, C. Johnbosco, A. Anspach, M. Meteling, J. Olijve, N. F. König, J. Leijten, Advanced Materials 2025, 37, 2501052

