## Supplementary information for "Vitamin C-Induced Photo-Redox Threshold Enables High-Fidelity Volumetric Printing of Pristine Collagen"

Supplementary Method 1

Supplementary Figures 1-11

Supplementary Tables 1-2

**Supplementary Method 1: Preparation of Biomaterials in PBS Buffer**

**Collagen resin:** Medical-grade purified type I collagen powder of bovine origin (CBPE2, Symatese) was dissolved in phosphate-buffered saline (PBS) to a final concentration of 5 mg/mL. The mixture was incubated at 4 °C for 24 hours to facilitate dissolution. To remove any undissolved material, the mixture was centrifuged (500 rpm, 5 min). Though the resin is kept on top of ice to suppress the self-assembly. The final solution appeared less transparent than collagen dissolved in water, likely due to self-assembly induced fibril formation. To complete the photoresin, a photoinitiator system comprising 0.2 mM ruthenium (Ru) and 2 mM sodium persulfate (SPS) (Advanced BioMatrix) was added to the solution. For formulations incorporating vitamin C (95210, Sigma-Aldrich), the required amount of vitamin C (different concentration as indicated) was added to the resin prior to the introduction of the Ru/SPS system.

**Fibrinogen resin:** Fibrinogen from bovine plasma (F8630, Sigma) was dissolved in PBS to a final concentration of 30 mg/mL. The mixture was incubated at 4 °C for 24 hours, yielding a completely clear solution. To complete the photoresin, a photoinitiator system comprising 0.2 mM Ru and 2 mM SPS was added to the solution. For formulations incorporating vitamin C, the required amount of vitamin C (different concentration as indicated) was added to the resin prior to the introduction of the Ru/SPS system.

**GelNB and GelSH resin:** Gelatin norbornene (GelNB) and thiolated gelatin (GelSH) were synthesized in-house based on a protocol adapted from our previous work. The GelNB and GelSH powders were co-dissolved in PBS to a final concentration of 20 mg/mL for each component (40 mg/mL total polymer concentration). To complete the photoresin, a 0.1 wt% photoinitiator lithium phenyl (2,4,6-trimethylbenzoyl) phosphinate (LAP) was added to the solution. For formulations incorporating vitamin C, the required amount of vitamin C (different concentration as indicated) was added to the resin prior to the introduction of the Ru/SPS system.

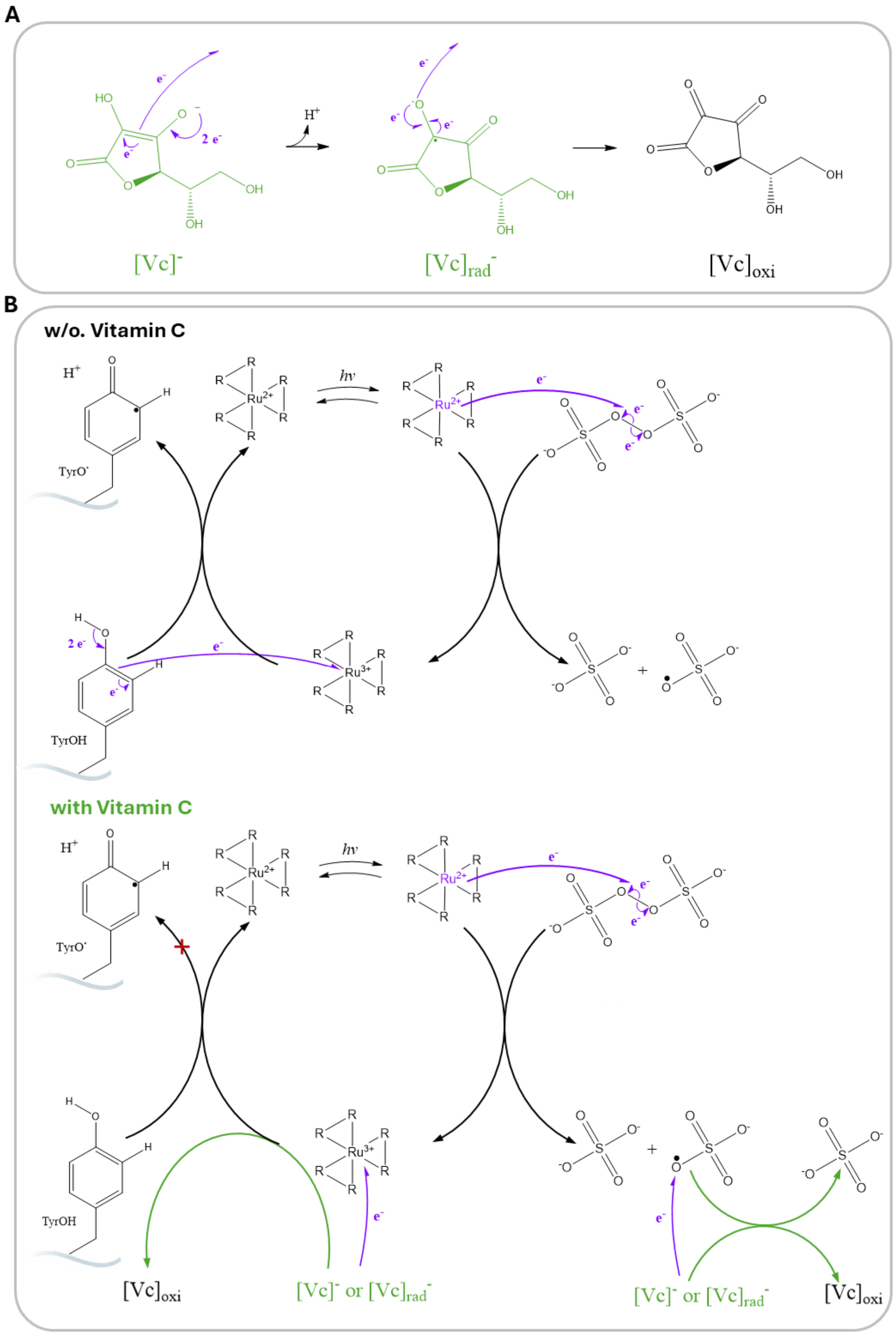


**Figure S1. Chemical structures and electron flow associated with the Ru/SPS redox system for dityrosine crosslinking, with and without vitamin C.** (A) Oxidation of vitamin C (ascorbic acid): The stepwise electron abstraction process is shown, where ascorbic acid ($[Vc]^-$) is oxidized via radical intermediates ($[Vc]_{rad}^-$) to its oxidized form ($[Vc]_{oxi}$), with curved arrows indicating the flow of electrons. (**B**) Schematic redox cycles during tyrosine crosslinking with (bottom) and without (top) vitamin C. In the absence of vitamin C (top), the Ru/SPS system generates persulfate radicals upon illumination (*hv*), which oxidize tyrosine residues to form tyrosyl radicals and subsequently dityrosine crosslinks. Electron transfer pathways are depicted in purple. In the presence of vitamin C (bottom), ascorbic acid serves as a sacrificial electron donor, reducing Ru(III) species back to Ru(II) and quenching sulfate radicals, thus modulating the redox cycle and providing control over the timing and extent of protein crosslinking. The roles of vitamin C in electron flow are highlighted in green.

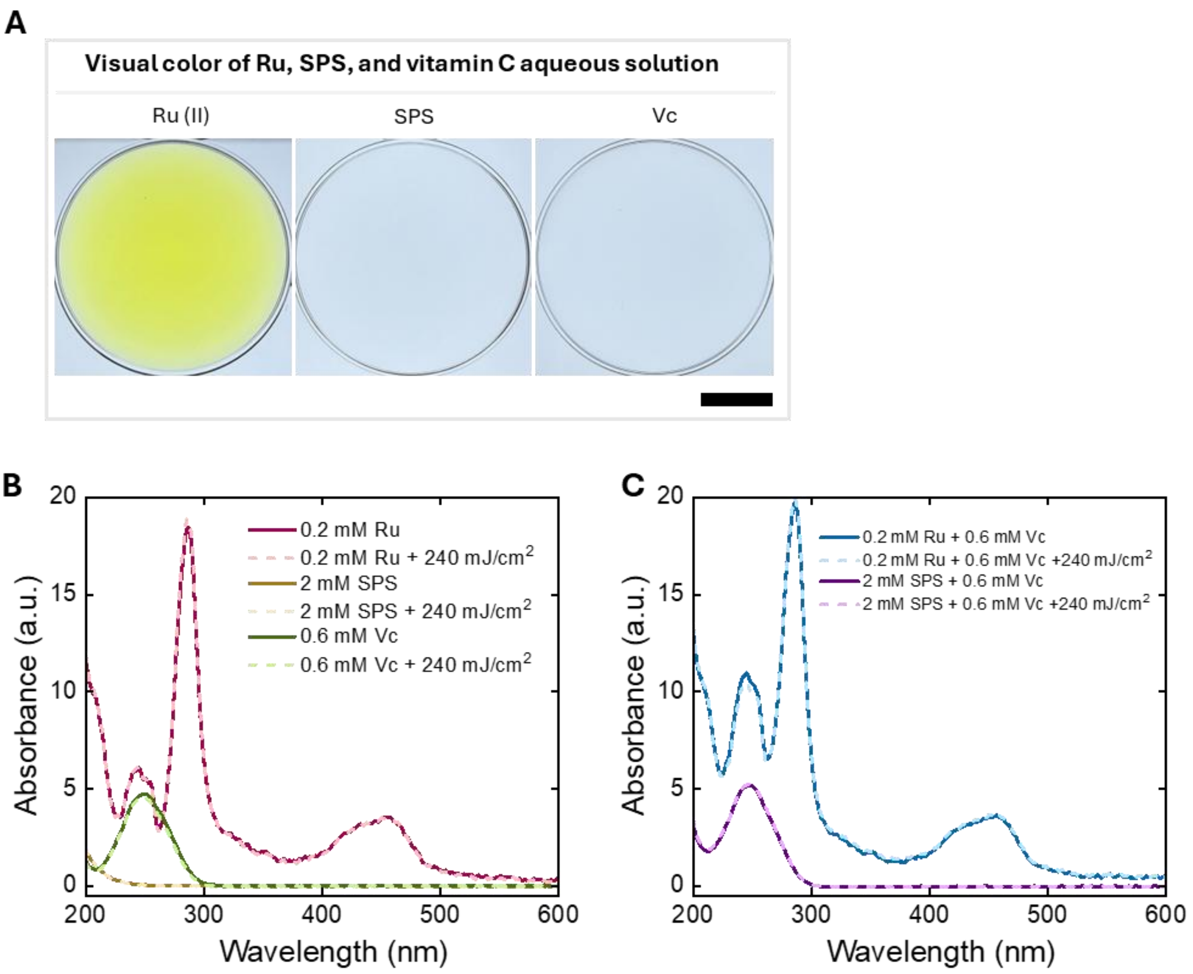


**Figure S2. Optical properties and absorption spectra of the photoresin components.** (**A**) Visual appearance of the individual aqueous solutions of Ru(II), SPS, and vitamin C (Vc), scale bar: 4mm. (**B**) UV-Vis absorption spectra of the individual components (0.2 mM Ru, 2 mM SPS, and 0.6 mM Vc) before (solid line) and after light exposure at a dose of 240 mJ/cm² (dash line). $Ru(bpy)_3^{2+}$ exhibited three main absorption peaks at 245 nm, 285 nm, and 455 nm. Vc displayed a single main absorption peak at 250 nm, whereas no absorption peaks were observed for SPS within the evaluated 200–700 nm wavelength range. (**C**) UV-Vis absorption spectra of the binary mixtures (0.2 mM Ru + 0.6 mM Vc, and 2 mM SPS + 0.6 mM Vc) before (solid line) and after light exposure (dash line) (240 mJ/cm²). Spectral comparisons indicated that no cross-reactions were observed between Ru and Vc, or between SPS and Vc, either in the absence of light or following illumination.

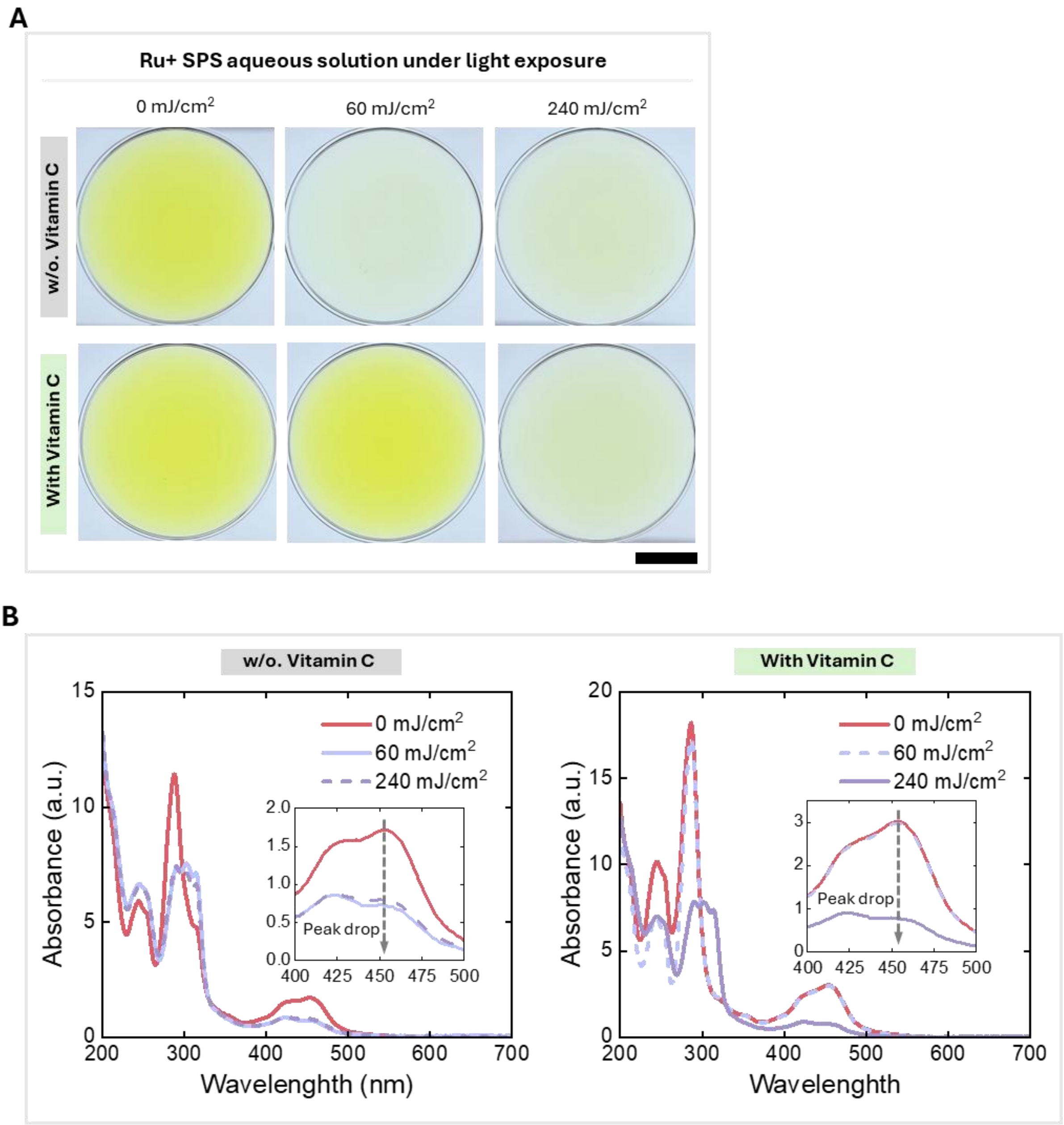


**Figure S3. Optical properties and photo-oxidation behavior of Ru/SPS aqueous solutions with and without vitamin C.** (**A**) Visual appearance of 0.2 mM Ru(II) / 2 mM SPS solutions without (top) and with (bottom) 0.6 mM vitamin C, following light exposures of 0, 60, and 240 mJ/cm$^2$. Scale bar: 4 mm. (**B**) Corresponding UV-Vis absorption spectra. Insets highlight the Ru(II) absorption peak at 455 nm. In the absence of vitamin C, the 455 nm peak droped significantly at 60 mJ/cm$^2$, indicating the rapid oxidation of Ru(II) to Ru(III), with no further change at 240 mJ/cm$^2$. In contrast, the presence of vitamin C suppressesd Ru(II) oxidation at 60 mJ/cm$^2$, preserving the 455 nm peak. At this dose, vitamin C was consumed preferentially by oxidation, evidenced by the decrease in its characteristic absorption peak near 250 nm. Once the vitamin C was exhausted at the higher exposure dose 240 mJ/cm$^2$, the oxidation of Ru(II) resumed, leading to a subsequent drop in the 455 nm peak. These results demonstrated that vitamin C effectively regulated the redox reaction and delayed the accumulation of Ru(III).

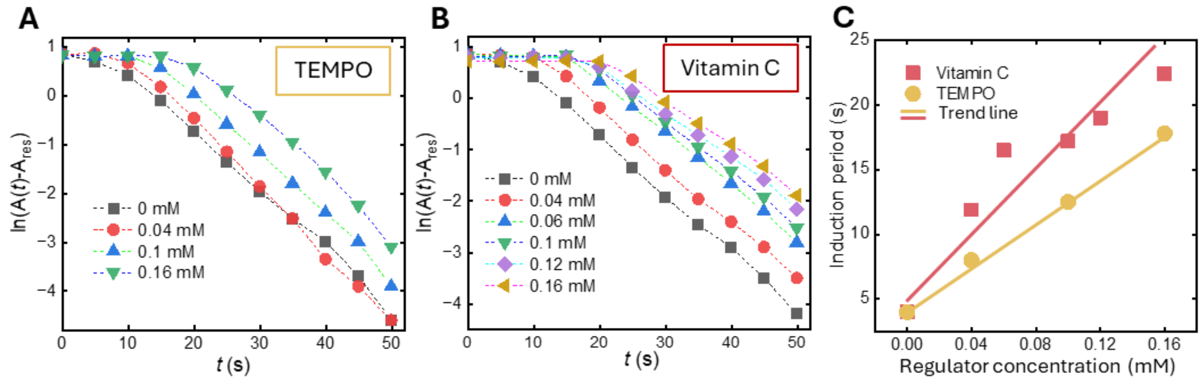


**Figure S4. Comparison of TEMPO and vitamin C suppression effect on Ru(II) photo-oxidation.** Time-resolved pseudo-first-order decay plots of the Ru(II) MLCT band monitored at 455 nm in the presence of varying concentrations of (**A**) TEMPO and (**B**) Vitamin C. The reaction speed was represented by ln(A(t)-Ares, where Ares is the residual absorbance at t = 55 s. (**C**) Linear dependence of the induction period on regulator concentration. Reactions were performed under continuous 405 nm light exposure 2 mW/cm$^2$) with 0.2 mM Ru and 2.0 mM SPS. Regulator concentrations (0-0.16 mM) were intentionally kept below the Ru concentration to evaluate their ability to suppress the reaction at sub-stoichiometric levels. In the absence of a regulator, the catalyst underwent rapid pseudo-first-order decay. Addition of either regulator slowed the decay and produced a concentration-dependent induction period where the Ru(II) signal remained essentially constant. These findings suggested the regulators acted as sacrificial reductants—consuming photogenerated oxidants Ru(III) and preventing net Ru(II) loss. vitamin C was markedly more effective than TEMPO, yielding substantially longer induction times at equivalent concentrations.

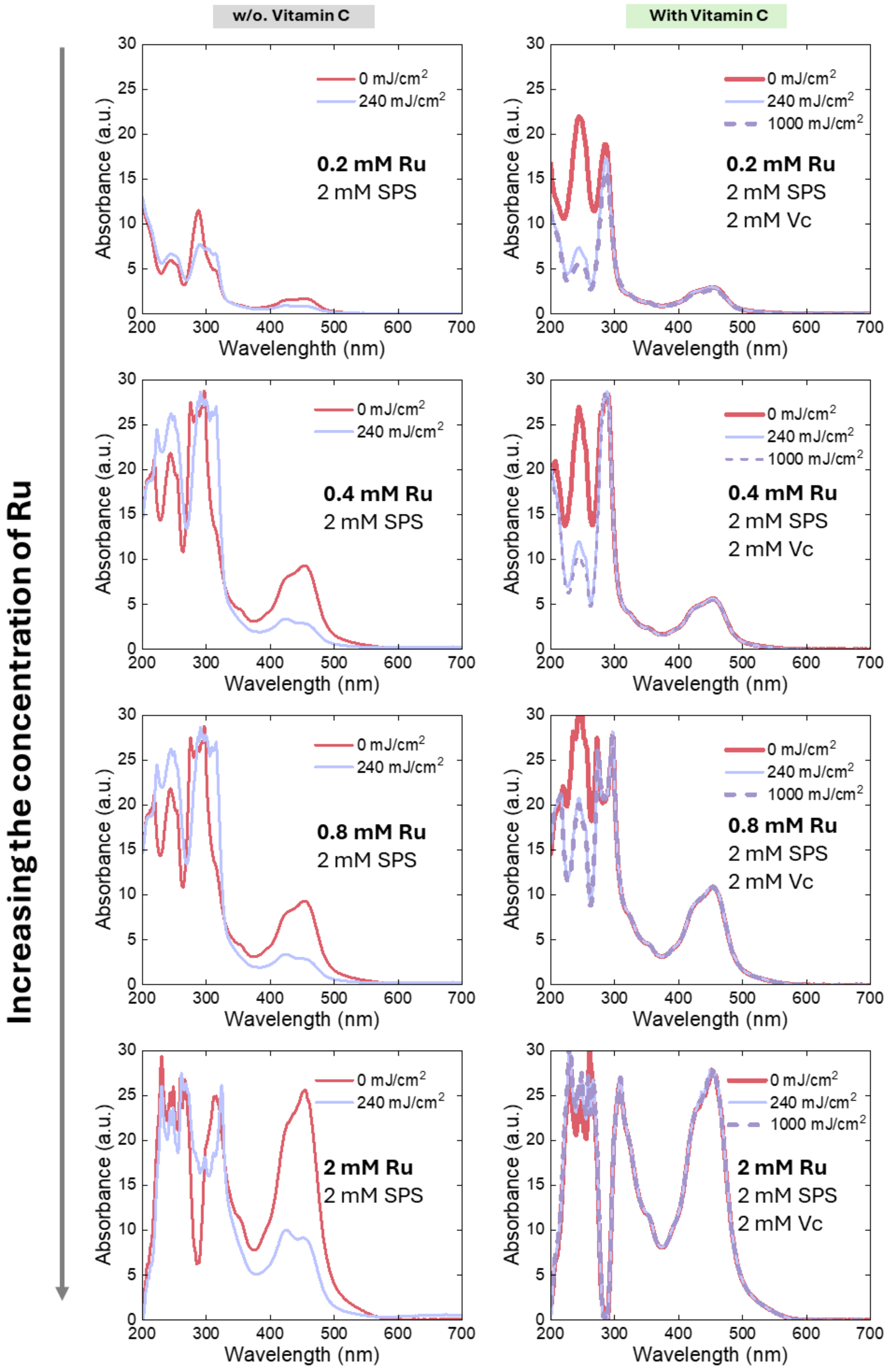


**Figure S5. Complete suppression of Ru(II) photo-oxidation by vitamin C at an equimolar concentration to SPS.** UV-Vis absorbance spectra of Ru/SPS aqueous solutions without (left column) and with (right column) vitamin C at varying Ru(II) concentrations (0.2, 0.4, 0.8, and

2.0 mM). The concentrations of SPS and Vc were fixed at 2 mM. In the absence of Vc, Ru(II) photo-oxidation occurs, evidenced by a drop in the absorbance peak at 455 nm across all Ru(II) concentrations after a light dose of 240 mJ/cm². In contrast, the addition of 2 mM Vc completely prevented this peak drop at 240 mJ/cm², and even at an elevated dose of 1000 mJ/cm². This demonstrated that Vc completely suppressed the redox reaction when its concentration is equal to that of SPS.

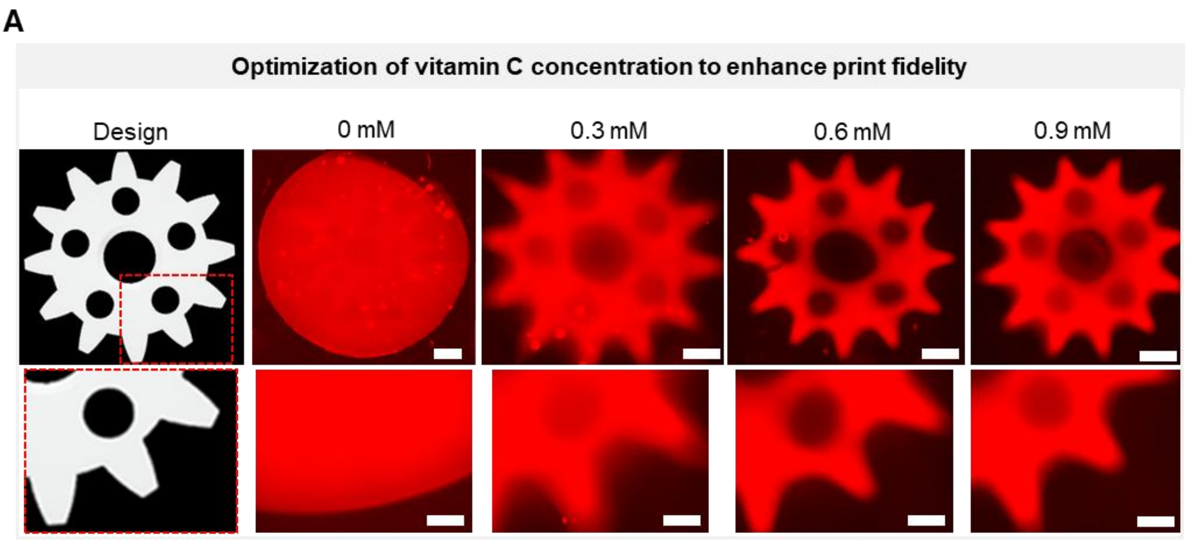


B

| 5 mg/ml collagen, 0.2 mM Ru / 2 mM SPS | | | | |
|---|---|---|---|---|
| **[Vc] (mM)** | 0 | 0.3 | 0.6 | 0.9 |
| **Light dose ($mJ/cm^2$)** | 67.8 | 300 | 650 | 1100 |
| **Print time (s)** | 6.8 | 30 | 65 | 110 |

**Figure S6. Optimization of vitamin C concentration for high-fidelity volumetric printing.** (A) Digital design of a gear model and corresponding fluorescence images of the printed collagen constructs formulated with varying concentrations of vitamin C (0, 0.3, 0.6, and 0.9 mM). The bottom row displayed magnified views of the gear teeth (red dashed boxes). demonstrating that without vitamin C, prints exhibited massive background curing even at the lowest allowable light doses, resulting in a complete loss of the intended geometry. The addition of 0.3 mM vitamin C partially mitigated this effect but was insufficient to fully suppress background curing, leaving the sharp features of the gear teeth blurred. In contrast, increasing the concentration to 0.6 mM and 0.9 mM effectively eliminated unintended background crosslinking. Both concentrations yielded high-fidelity constructs with equally well-resolved structural details. Scale bars: Top panel: 1 mm; bottom panel: 2 mm. (B) Summary table detailing the corresponding light dose and required print time for each vitamin C concentration for gear print using a formulation of 5 mg/mL collagen, 0.2 mM Ru, and 2 mM SPS.

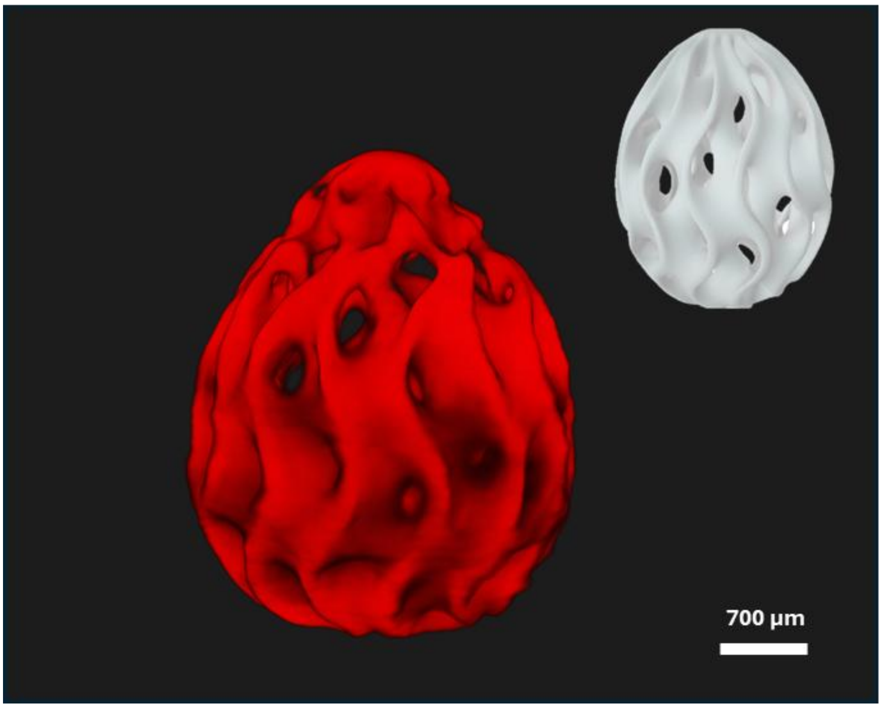


**Figure S7. High-fidelity volumetric printing of a collagen-based gyroid structure.** Light sheet microscopy imaging of an eggshell gyroid design fabricated with vitamin C supplemented collagen resin. The complex porous architecture was accurately reproduced. Inset: original CAD model. Scale bar: 700 µm.

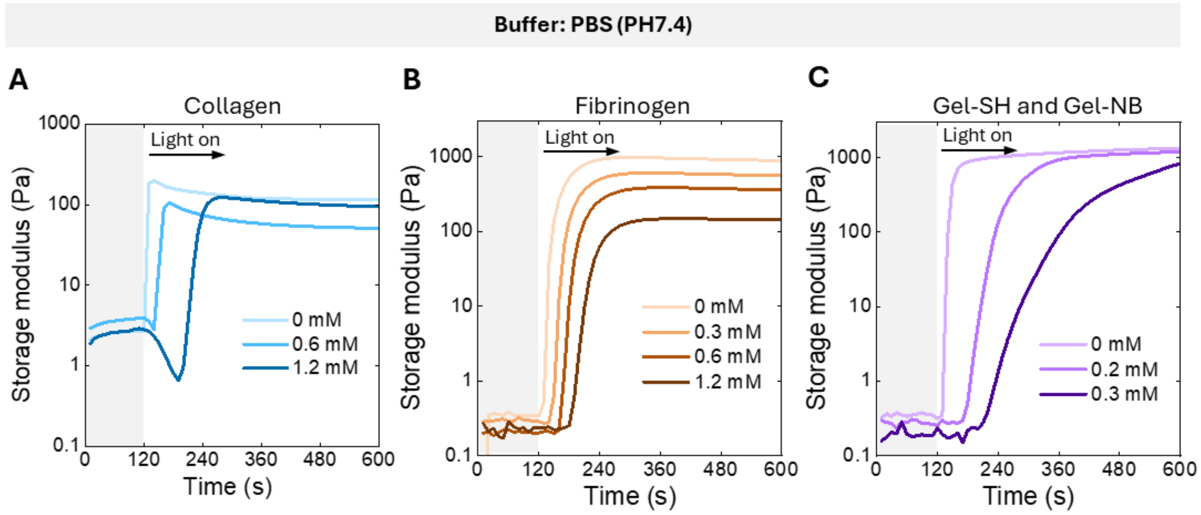


**Figure S8. Universal applicability of vitamin C introducing inhibitory control in step-growth polymerization across diverse biomaterials.** Time-sweep rheology showing the storage modulus of various biomaterials dissolved in PBS. Collagen and fibrinogen formulations utilized a photoinitiator system of 0.2 mM Ru(II) and 2 mM SPS, with varying concentrations of vitamin C acting as the redox regulator. While Gel-SH and Gel-NB used 0.1 wt% LAP as initiator, with vitamin C added as polymerization inhibitor. Light irradiation commences at 120 s. Across all materials, vitamin C introduced a tunable, concentration-dependent induction period, thereby delaying the onset of crosslinking. (**A**) 5 mg/mL collagen. (**B**) 30 mg/mL fibrinogen. (**C**) 20 mg/mL Gel-SH mixed with 20 mg/mL Gel-NB. The consistent induction behavior across different macromolecular systems demonstrated the broad applicability of vitamin C in introducing inhibitory control in step-growth polymerization.

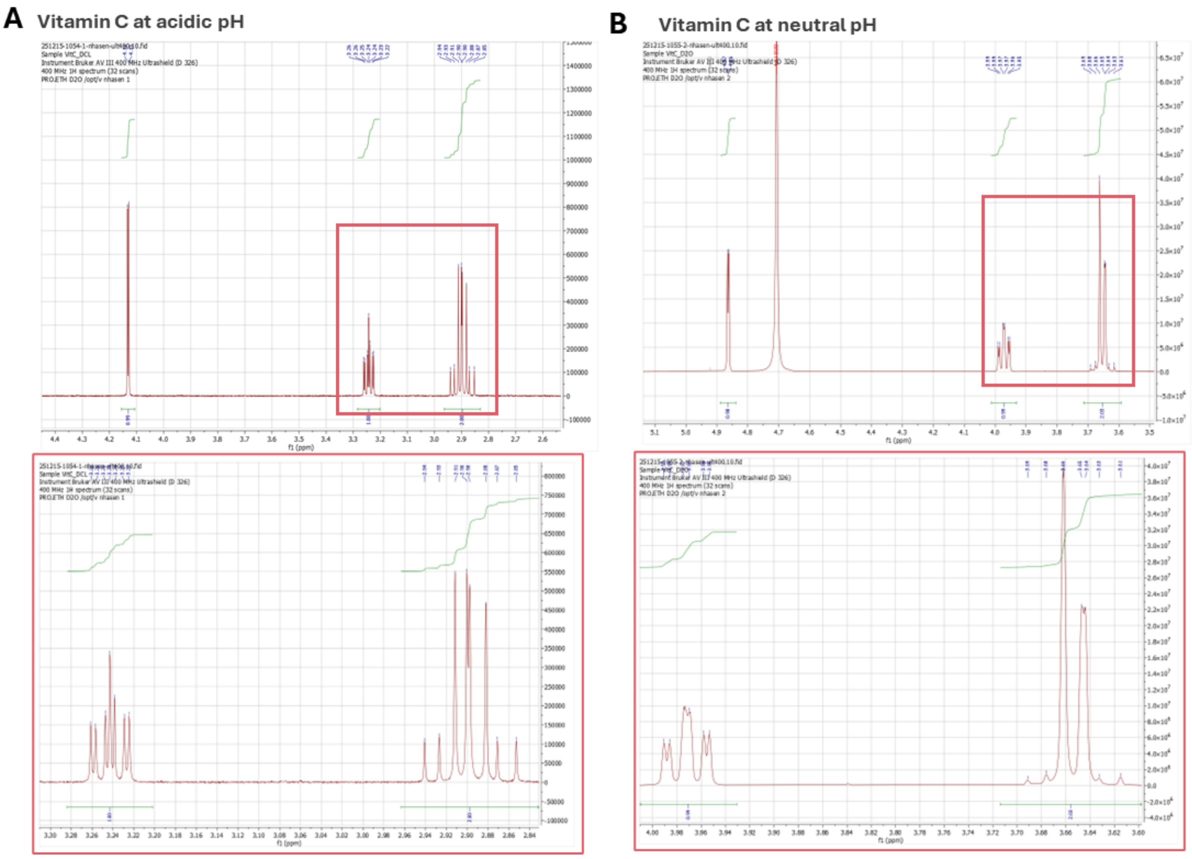


**Figure S9. H NMR of vitamin C under acidic and neutral conditions.** $^{1}$H NMR spectra of vitamin C (L-ascorbic acid) under (**A**) acidic conditions in deuterium oxide ($D_2O$, matching the collagen printing formulation in water, PH 3.5) and under (**B**) near-neutral conditions in deuterium chloride (DCl, collagen in neutral-buffer condition, pH 7). The spectra showed no appearance of new peaks and no loss of the characteristic vitamin C signals across conditions, indicating that vitamin C remained stable between acidic and neutral environments (red boxes and zoom-ins at the bottom show the peaks of interest).

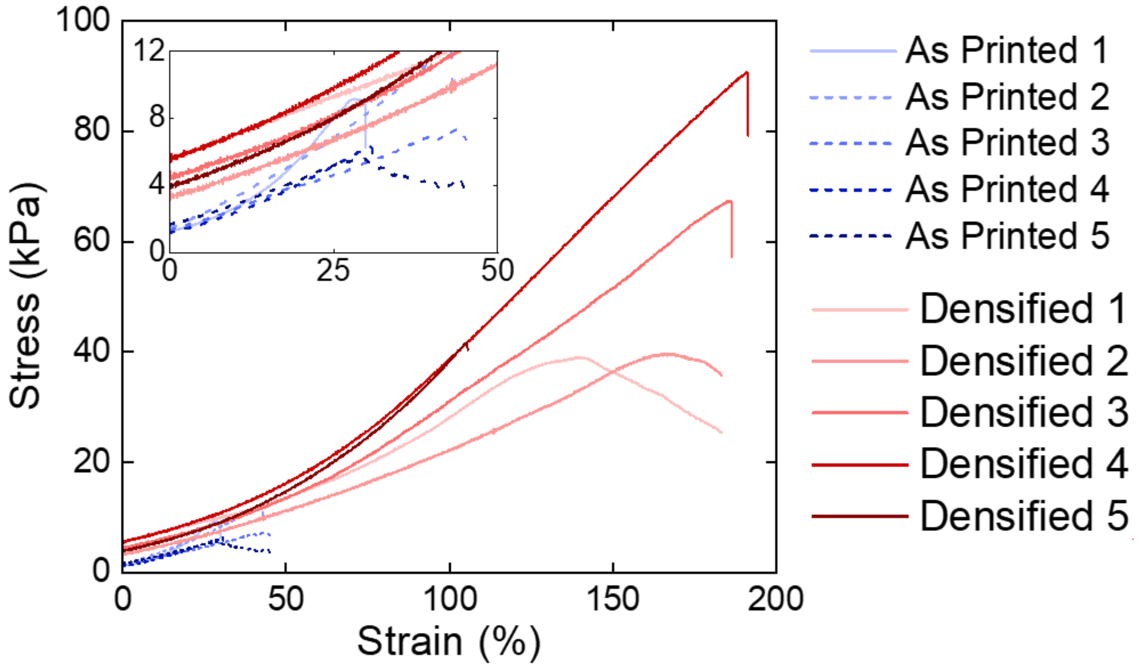


**Figure S10. Tensile stress-strain curves of VP printed "dog-bones" specimens.** Five replicate samples ($n = 5$) were tested for both the as-printed (blue lines) and densified (red lines) groups. The densified specimens demonstrated significantly enhanced mechanical properties, achieving higher ultimate tensile stress and overall elongation. The inset magnified the low-strain region (0–50%) to clearly detail the early yielding and failure behavior of the as-printed samples.

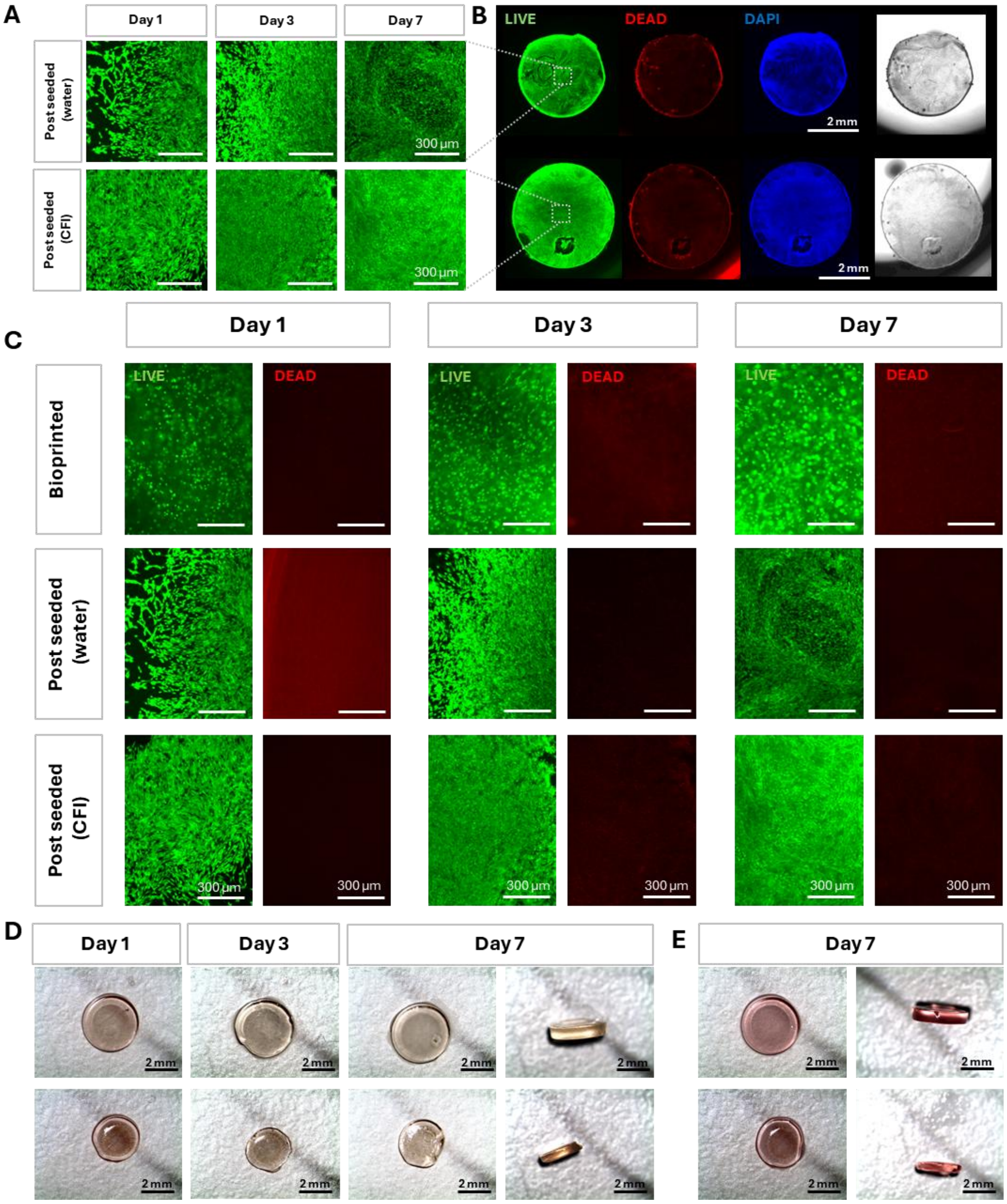


**Figure S11. Viability and structural integrity of 3T3-seeded collagen prints over 7 days.** Collagen disks were fabricated by volumetric printing using RU-SPS with vitamin C supplementation (0.6 mM) and evaluated after seeding with 3T3 fibroblasts. Cell viability was assessed by Live/Dead staining (LIVE, green; DEAD, red; nuclei counterstain DAPI, blue) at days 1, 3, and 7. **(A)** Representative fluorescent images of Live signal over time for post-seeded constructs printed from collagen resuspended in in water (top row) or glucose rich CFI buffer (bottom row). In CFI buffer, collagen densification post printing is not observed. **(B)** Overview of the printed disks at day 7 showing LIVE (green), DEAD (red), and DAPI (blue) channels, alongside corresponding brightfield views, printed form collagen resuspended in water (top) or CFI buffer (bottom). **(C)** Additional live/dead imaging at days 1, 3, and 7 for three conditions: bioprinted (cell-laden printing), post-seeded on printed collagen disk with collagen resuspended in water and CFI buffer respectively. **(D)** Brightfield overviews of printed disks seeded with cells over 1 week (top views and side views) demonstrating construct integrity and material stability, with no macroscopic degradation observed after culture. **(E)** Acellular controls for after 1 week.

**Supplementary Table 1: Measurement of ambient light irradiance at selected wavelengths.** Irradiance values were recorded using a calibrated photodiode power sensor (S120VC, Thorlabs) with a spectral response range of 200–1100 nm. The measurements reflected ambient illumination from LED tube lights commonly used in the laboratory environment.

| Wavelength (nm) | Irradiance ($mW/cm^2$) |
| --- | --- |
| 405 | 0.28 |
| 450 | 0.22 |
| 455 | 0.22 |

**Supplementary Table 2: Measurements of printed cylinder dimensions before and after densification, and corresponding shrinkage ratios.** Lateral and vertical dimensions (in millimeters) were reported for each sample in both As-printed and Densified states.

| Cylinder samples | Measured dimentions (mm) | | | | Shrinkage ratio (%) | |
|---|---|---|---|---|---|---|
| | As printed | | Densified | | | |
| | Lateral | Vertical | Lateral | Vertical | Lateral | Vertical |
| 1 | 8.6 | 4.4 | 4.1 | 2.1 | 52.3 | 52.3 |
| 2 | 8.6 | 4.4 | 4.1 | 2.1 | 52.3 | 52.3 |
| 3 | 8.6 | 4.4 | 4.2 | 2.1 | 51.2 | 52.3 |
| 4 | 8.6 | 4.4 | 4.1 | 2.1 | 52.3 | 52.3 |
